# NOD-like receptor genes evolve under diversity-enhancing mechanisms in a fungal species complex

**DOI:** 10.1101/2025.09.29.679196

**Authors:** S. Lorena Ament-Velásquez, Sven J. Saupe

## Abstract

Fungi harbor diverse arrays of genes encoding NOD-like receptors (NLRs), key intracellular immune proteins found in plants, animals, and bacteria. Some fungal NLRs are known to control regulated cell death (RCD) in the context of allorecognition, the capacity to recognize conspecific nonself. However, the function of most fungal NLR genes remains unknown. Here, we characterize the evolution of the NLR repertoire in the *Podospora anserina* species complex. We show that the vast majority of the NLRs display effector domains known to be involved in RCD execution. Moreover, NLRs undergo more rapid gene turnover than random genes, show higher d_N_/d_S_ values, and faster evolutionary rates. A subgroup of NLRs, distinguished by superstructure-forming repeats with very high sequence identity (high internal conservation, HIC), evolved independently multiple times. We found that HIC NLRs are more associated with transposable elements, exhibit higher nucleotide diversity partially driven by repeat-induced point mutation (RIP), and show elevated Tajima’s *D* values indicative of balancing selection. Furthermore, HIC NLR phylogenies do not recapitulate species relationships, which we determined is caused by both balancing selection and introgression. In addition, we identified cases of repeat exchange between distinct HIC NLR genes, implying that novel binding specificities may evolve through repeat shuffling, thereby increasing allelic diversity. Finally, we determined that NLR-like genes with HIC repeats exist outside of the fungal realm, arguing for similar dynamics in other taxa. Overall, these findings suggest that fungal NLRs evolve under diversity-enhancing mechanisms and display selective signatures consistent with a general immune function.

## Introduction

Genes involved in self-nonself recognition are typically subject to evolutionary regimens that diverge from the combination of purifying selection and genetic drift governing much of the genome (Ebert and Fields 2020; Moya et al. 2025). In immunity-related loci, the persistent arms race with pathogens leads to rapid diversification and frequent allele turnover, driven by positive (diversifying) selection and birth-and-death evolutionary dynamics (Nei and Rooney 2005). Such loci might also display high levels of polymorphism, maintained by various forms of balancing selection, including overdominance, negative frequency-dependent selection, and spatiotemporal variation in pathogen pressures (Woolhouse et al. 2002; Tellier et al. 2014). Loci controlling somatic or vegetative rejection between tissues (i.e., allorecognition) typically also evolve under balancing selection (Nydam and DeTomaso 2011). Thus, patterns of selection and polymorphism across the genome can be used to identify novel genetic systems involved in biotic interactions of understudied organisms (Kaur et al. 2020; Zhao et al. 2015).

Nucleotide-binding oligomerization domain (NOD)-like receptors (NLRs) stand out as a prominent class of immune receptors across the Tree of Life (Jones et al. 2016). These intracellular proteins mediate both antiviral or antibacterial innate immunity in mammals (Sundaram et al. 2024), represent the largest group of disease resistance (*R*) genes in plants (Guo et al. 2025), and contribute to antiphage defense in bacteria (Gao et al. 2022; Kibby et al. 2023). Notably, NLRs do not represent a single conserved protein lineage maintained over deep evolutionary time. Instead, they are characterized by a shared domain architecture composed of ancient, immune-related modules that have convergently reassembled multiple times in the context of self-nonself recognition and pathogen defense (Urbach and Ausubel 2017). Specifically, an NLR gene has a tripartite organization, consisting of an N-terminal effector domain that usually triggers regulated cell death (RCD), a central NOD module, and a C-terminal sensor domain (Jones et al. 2016). NLR activation involves ligand-induced oligomerization, where binding to the sensor domain triggers assembly into a wheel-shaped complex and subsequent activation of the effector domain (Hu and Chai 2023). The NOD module is a P-loop ATPase from the STAND superfamily that mostly belongs to either one of two major lineages: NACHT (named after the NAIP, CIIA, HET-E, and TLP1 proteins) or NB-ARC (Nucleotide-Binding adaptor shared by APAF1, plant R proteins, and CED4) (Jones et al. 2016). Different domains can act as an N-terminal effector domain, including members of the death-fold superfamily in animals, and the Toll-interleukin receptor (TIR) or the membrane-targeting coiled-coil domains in plants. The C-terminal sensor domain is composed of superstructure-forming tandem repeats, typically leucine-rich repeats (LRRs) in mammals and plants, while other types of superstructure forming repeats such as WD40, Ankyrin (ANK) or tetratricopeptide (TRP) repeats are more widespread in other taxonomical groups (Lange et al. 2011; Teng et al. 2023; Urbach and Ausubel 2017; Koutsouveli et al. 2024).

Filamentous fungi display large NLR repertoires, but their function has so far only been studied in relation to allorecognition (Daskalov et al. 2020; Wojciechowski et al. 2022; Dyrka et al. 2014). Allorecognition is a general immune challenge in modular organisms capable of forming somatic chimeras such as sessile marine invertebrates and protochordates, slime molds, and myxobacteria (Daskalov et al. 2017; Kaimer et al. 2023; Kundert and Shaulsky 2019; Lakkis et al. 2008; Nicotra 2019). In fungi, this challenge is particularly pressing because somatic cell fusion events between different individuals occur spontaneously, resulting in the mixing of cytoplasmic contents, sometimes including nuclei. Mycoviruses are essentially transmitted via this route between individuals (Kotta-Loizou 2021). In most cases, however, somatic fusion is interrupted by an incompatibility reaction leading to RCD of the fused cells, which prevents further cytoplasmic exchange and limits viral transmission (Biella et al. 2002; Debets et al. 2012; Zhang et al. 2014). Incompatibility is controlled by a series of unlinked polymorphic loci termed *het* or *vic*, usually between six and 11 per species (Esser 2006; Auxier et al. 2024; Perkins 1988; Arshed et al. 2023). Since a single genetic difference between strains at any of those loci is sufficient to trigger incompatibility, natural populations are split into tens to hundreds of compatibility groups. As commonly found in other allorecognition loci, *het* genes display signatures of balancing selection like equilibrated allele frequencies and trans-species polymorphism (Auxier et al. 2024; Zhao et al. 2015; Milgroom et al. 2018; Ament-Velásquez et al. 2022). Importantly, many of the identified *het* genes across species encode NLRs or NLR-like proteins (Chevanne et al. 2010; Auxier et al. 2024; Heller et al. 2018; Choi et al. 2012; Arshed et al. 2023). These *het* NLRs trigger RCD upon recognition of their specific, incompatible ligand: the cognate determinant from a different strain (Ament-Velásquez et al. 2022; Heller et al. 2018; Saupe et al. 1995). It can thus be stated that NLRs can have a role in immune defense and control of RCD in fungi, at least in the context of allorecognition. Yet, there are many more NLRs (on average 57 per genome) than *het* loci in a given fungal species (Dyrka et al. 2014). So what functions do the remaining NLRs serve?

Considering their prominent role in immune defense in other taxonomic groups, it has been proposed that the bulk of fungal NLRs play general roles in defense and symbiosis establishment, while just a few of them have been co-opted to function in allorecognition (Paoletti and Saupe 2009; Daskalov 2023, 2025). Under this model, detection of positive and balancing selection in NLRs in general would support their role in pathogen defense and immunity. By contrast, evolutionary signatures consistent with background genomic evolution would be more indicative of conserved, housekeeping functions, such as nutrient response or development. Comparative genomic studies have frequently revealed lineage-specific expansions and contractions of NLR gene families, along with instances of diversifying selection (Iotti et al. 2012; Van der Nest et al. 2014; Zhao et al. 2015; Dauphin et al. 2021; Dyrka et al. 2014; Kubicek et al. 2019; Armaleo et al. 2019). However, with few exceptions, these findings emerged from broad comparative genomic analyses rather than studies specifically focused on NLRs. Moreover, neither NLRs nor *het* genes are generally conserved across species, limiting the utility of cross-species comparisons as phylogenetic distance increases. To better understand the evolution and broader functional roles of this gene class — beyond their established involvement in allorecognition — fine-scale evolutionary analyses dedicated specifically to fungal NLRs are now required.

The saprotrophic ascomycete *Podospora anserina* is a major model for the study of allorecognition, with its full complement of *het* genes characterized (Silar 2020; Clavé et al. 2024). Importantly, out of its nine *het* loci, four involve NLRs directly: *het-z*, *het-d*, *het-e*, and *het-r*. Specifically, the *het-z* locus encodes an NB-ARC+TPR NLR with a Patatin lipase domain as N-terminal effector. The other three, *het-d*, *het-e*, and *het-r*, are closely related but unlinked NLRs encoding HNWD proteins named after their constitutive parts: an N-terminal <u>H</u>ET domain, a <u>N</u>ACHT central domain, and <u>WD</u>40 C-terminal repeats. The HET module is a cell-death-inducing domain distantly related to TIR, which is very frequent in fungi and found in several other non-NLR *het* genes (Dyrka et al. 2014; Zhang et al. 2014; Zhao et al. 2015; Clavé et al. 2024). The HNWD gene family, in turn, is part of a larger collection of NLRs called NWD genes that have NACHT and WD40 domains but alternative N-terminal domains (Paoletti et al. 2007; Daskalov et al. 2012). A member of this family (*nwd2*) is also indirectly involved in incompatibility, as part of a two-component RCD system controlled by amyloid signaling (Daskalov et al. 2012). Beyond these genes, *P. anserina* harbors many more NLR genes of unknown function.

The WD40 domain that defines ligand-specificity in HNWD genes is very peculiar in the sense that individual repeats are nearly identical to each other (80-100% identity), a condition termed high internal conservation or HIC (Saupe et al. 1995; Dyrka et al. 2014). Paradoxically, while the repeats are nearly identical, four key amino acid positions are hypervariable and under diversifying selection, likely controlling the ligand-specificity of the WD40 sensor domain (Paoletti et al. 2007; Chevanne et al. 2010; Ament-Velásquez et al. 2025). These HIC WD40 domains are polymorphic in wild populations and have a tendency to lose, gain, or shuffle repeats. The exact molecular mechanism maintaining HIC is unknown, but it is hypothesized that unequal crossing-overs between misaligned repeats drive concerted evolution within individual genes (Saupe et al. 1995). Different lines of evidence suggest that even intergenic repeat exchanges occur between members of the NWD family, possibly by ectopic gene conversion (Paoletti et al. 2007; Ohta 2010; Ament-Velásquez et al. 2025). First, pseudogenized NWD genes contain abundant C-to-T mutations, a signature of a fungal genome defense mechanism known as Repeat-Induced Point mutation (RIP). This system targets repeated sequences based on high similarity, including transposable elements (TEs), as well as newly duplicated genes (Hamann et al. 2000; Gladyshev and Kleckner 2014). However, the WD40 repeats of the NWD pseudogenes are nearly intact, as if rescued by the HIC process (Paoletti et al. 2007). Second, completely identical individual repeats can be found within repeat arrays of divergent members of the gene family (Ament-Velásquez et al. 2025), implying that intergenic repeat exchange can further increase diversity. Importantly, HIC is not unique to the HNWD genes or to *P. anserina.* Around 13% of NLRs in Dikarya display HIC in WD40, ANK, or TPR repeats (Dyrka et al. 2014), making the distinction of NLRs into HIC and non-HIC classes of general relevance.

The genomic resources available for *P. anserina* and six closely related species provide a unique opportunity to investigate NLR evolution within a functionally-informed framework and at a relevant phylogenetic depth (Ament-Velásquez et al. 2024). In this study, we undertake an analysis of NACHT and NB-ARC NLRs phylogenetically related to *het* genes. We first establish a detailed characterization of NLR architectures and their relationships within *P. anserina*, before assessing their fate in other *Podospora* species. We then explore their genomic context and population diversity, while making a distinction between NLRs with and without HIC. We attempt to disentangle individual instances of balancing selection from potential introgression with other species. We develop a collection of individual TPR repeats of related HIC NLRs in the search for intergenic repeat exchanges. And finally, we establish that there is also an association between HIC repeats and the NLR protein architecture outside of the fungal realm. Altogether, this study provides a framework for understanding the evolutionary dynamics of fungal NLRs in the context of allorecognition and beyond.

## Results

### NLRs in *P. anserina* show domain reshuffling, effector domains associated to regulated cell death, and independent instances of high internal conservation

To find and characterize the proteins related to functionally characterized RCD-inducing NLRs, we focused on their NACHT and NB-ARC ATPase domains (Leipe et al. 2004). We chose the NACHT domain of the HET-E and NWD4 proteins and the NB-ARC domain of the PaPLP1 protein (part of *het-z*) as representatives and performed BLASTp and tBLASTn searches on the reference genome of *P. anserina*, strain S+ (Espagne et al. 2008). We retained the BLAST hits that overlapped a gene model in the reference annotation and that had a recognizable NACHT and NB-ARC domains, defined by having a P-loop (Walker A) motif, composed of the sequence GxxxxGK[ST], and a Walker B motif with a conserved aspartate (Koonin and Aravind 2000; Walker et al. 1982; van der Biezen and Jones 1998).

Automatic annotation of NLR gene models can be of low quality due to their repetitive C-termini and uncharacterized domains (Tørresen et al. 2019; Yuen et al. 2014). Indeed, comparisons with alternative annotations of the same reference genome (Lelandais et al. 2022; Ament-Velásquez et al. 2024) revealed inconsistencies between gene models. Thus, we performed manual curation of all selected genes using the splicing patterns of RNAseq data from *P. anserina* and *Podospora comata* (Vogan et al. 2019), BLASTp searches in NCBI, and descriptions of known domains (**Table S1**). After gene-model corrections, we kept a total of 54 sequences for downstream analysis, although the manual curation process indicated that several of these genes are potential pseudogenes affected by stop codons, frame-shifts, and TE insertions (**Figure 1** and **2**). Conversely, inspection of the same genes in other *P. anserina* strains with long-read, high-quality assemblies (Vogan et al. 2019, 2021) showed that some of these pseudogenized genes are sometimes intact depending on the strain. In addition, we detected at least 12 more pseudogenized remains of NWD relatives along the S+ genome that did not fulfill our NACHT minimum requirements. We failed to identify the N-or C-terminal domains of some genes, many of them pseudogenized at various degrees (**Table S2**). Most of the NACHT sequences (40) in our dataset are of the NAIP-like family (Leipe et al. 2004; Bonometti et al. 2025). Hence, as an outgroup we used the *P. anserina* gene *Pa_6_7270*, which belongs to the TLP1-like family and is closest to our NLR set within the *P. anserina* genome. Below we refer to the 41 NACHT genes in our dataset as the NWD-related NLRs. The remaining 13 genes have an NB-ARC domain and lack an obvious outgroup.

**Figure 1.**
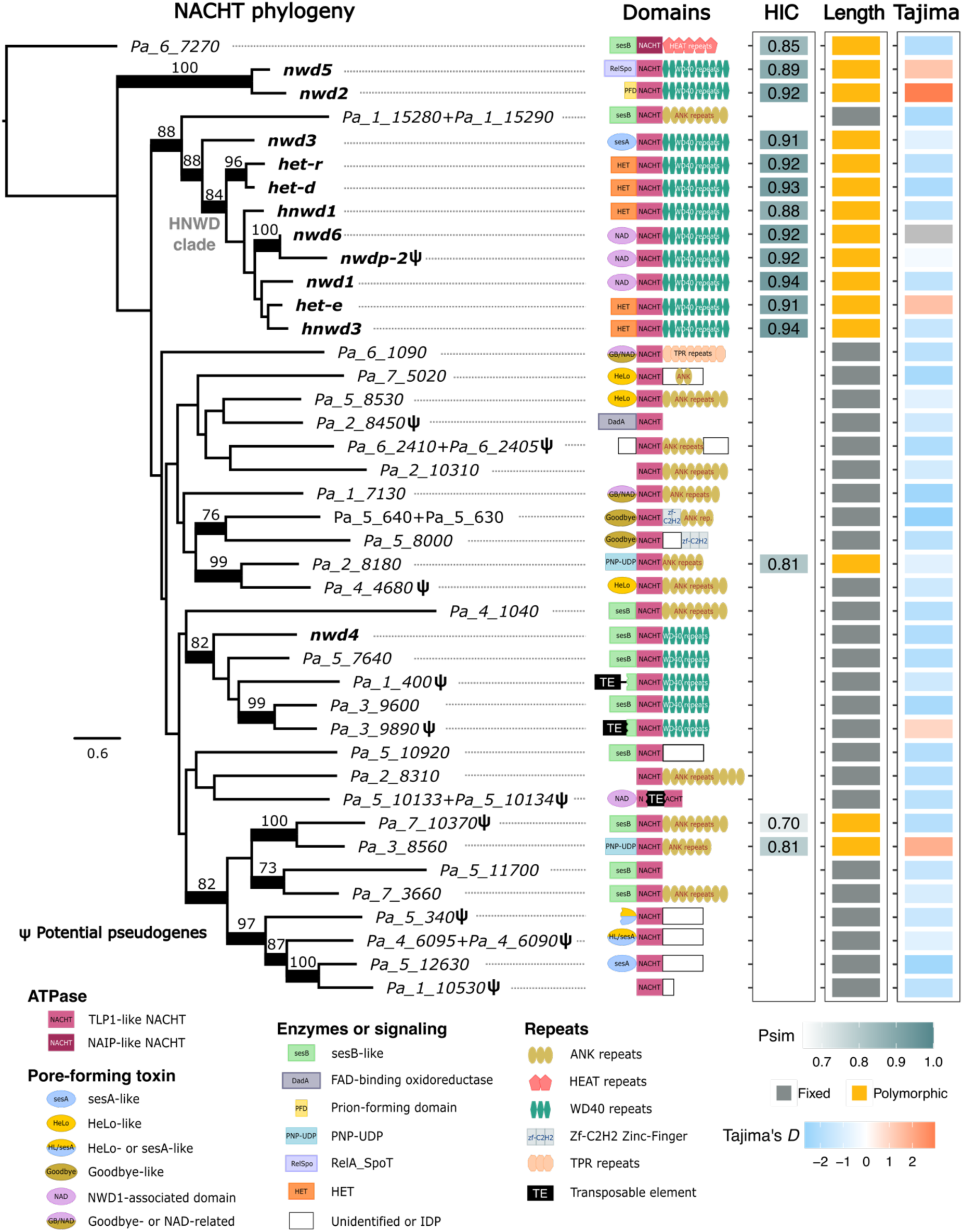
Phylogenetic relationships and main features of NWD-related genes in *P. anserina*. To the left, a maximum likelihood phylogeny of the core NACHT domain is drawn with branch lengths proportional to the scale bar (amino acid substitutions per site). Branches with nonparametric bootstrap support values above 70 are thickened; lower bootstrap values are omitted. For each gene, a diagram of the domain architecture is shown (not at scale). Genes previously classified as NWD genes are highlighted in bold. To the right, three different gene traits are depicted in colored heat maps. A gene is considered to have high internal conservation (HIC) if the similarity coefficient of its repeats, *Psim*, is equal to or larger than 0.7. A given gene might display allele length polymorphism based on the number of repeats (Polymorphic) in the long-read assemblies, or all alleles might be of equal length (Fixed). The average Tajima’s *D* statistic of the 10 kb-long overlapping windows (1 kb steps) that overlap with each gene is shown to the far right (data from the Wageningen Collection; Ament-Velásquez et al. 2022). Some gene models were merged into a single gene after manual curation (e.g., *Pa_4_6095+Pa_4_6090*). IDP: intrinsically disordered protein. **Figure 1 ALT TEXT.** Phylogeny of the NACHT NLR genes with different traits per gene marked to the right. The legend below the phylogeny illustrates the different kinds of protein domains in the NLR genes.

**Figure 2.**
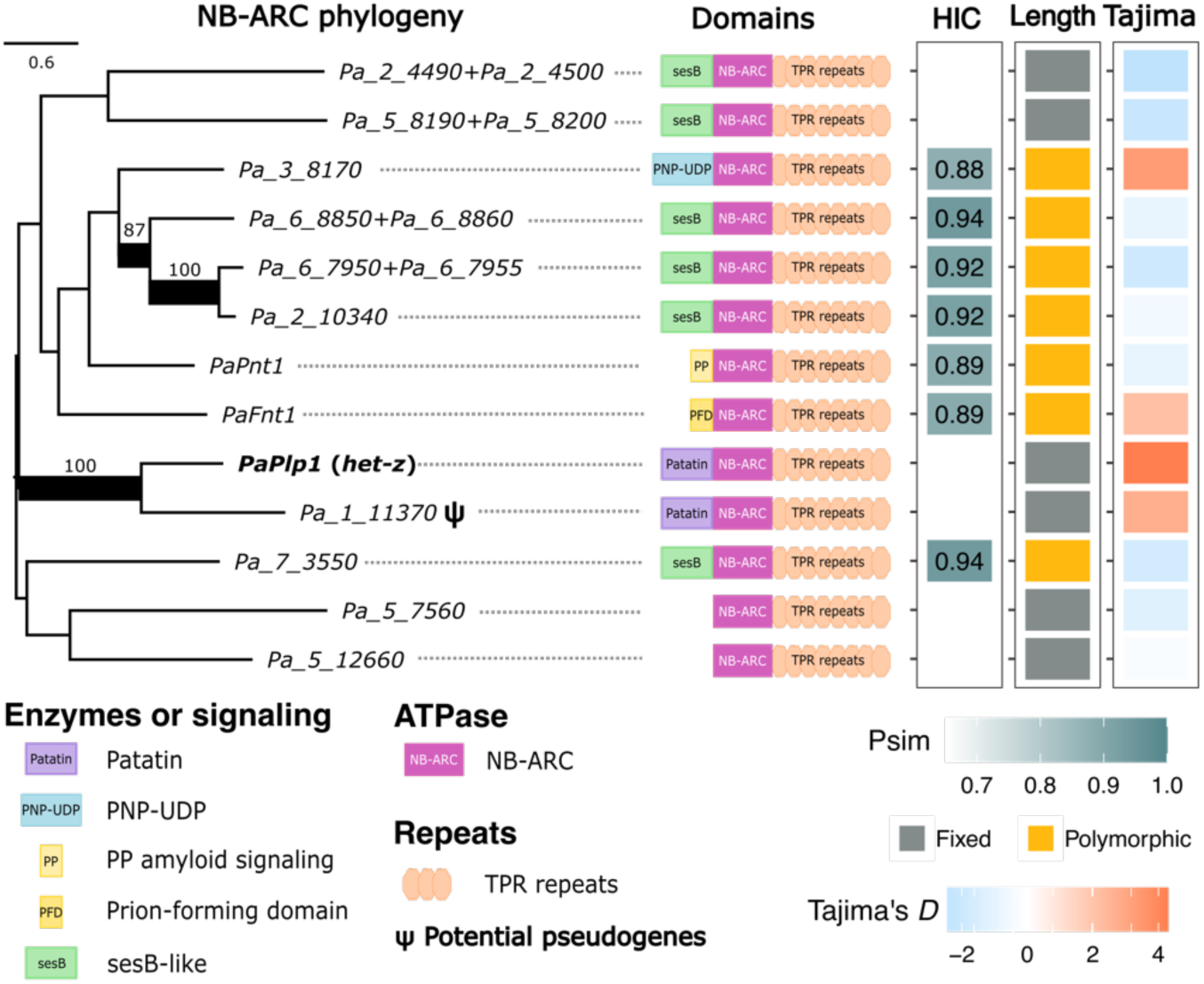
Phylogenetic relationships and main features of NB-ARC NLR genes in *P. anserina*. To the left, a maximum likelihood phylogeny of the core NB-ARC domain is drawn with branch lengths proportional to the scale bar (amino acid substitutions per site), rooted with the midpoint method. Branches with nonparametric bootstrap support values above 70 are thickened; lower bootstrap values are omitted. For each gene, a diagram of the domain architecture is shown (not at scale). The *PaPlp1* gene known to have an allorecognition function (*het-z*) is highlighted in bold. To the right, three different gene traits are depicted in colored heat maps. A gene is considered to have high internal conservation (HIC) if the similarity coefficient of its repeats, *Psim*, is equal to or larger than 0.7. A given gene might display allele length polymorphism based on the number of repeats (Polymorphic) in the long-read assemblies, or all alleles might be of equal length (Fixed). The average Tajima’s *D* statistic of the 10 kb-long overlapping windows (1 kb steps) that overlap with each gene is shown to the far right (data from the Wageningen Collection; Ament-Velásquez et al. 2022). Some gene models were merged into a single gene after manual curation (e.g., *Pa_6_8850+Pa_6_8860*). **Figure 2 ALT TEXT.** Phylogeny of the NB-ARC NLR genes with different traits per gene marked to the right. The legend below the phylogeny illustrates the different kinds of protein domains in the NLR genes.

We aligned the core sections of either the NACHT (Koonin and Aravind 2000) or NB-ARC (van der Biezen and Jones 1998) domains to infer maximum likelihood phylogenies (Figure 1 and **2**). The NLRs in *P. anserina* are highly divergent, which reduces the phylogenetic signal in their alignments. This high divergence is in part due to deep phylogenetic relationships. For example, homologs of both *nwd2* and *het-e* are abundant in Ascomycete and Basidiomycete genomes (Van der Nest et al. 2014), suggesting their split goes back to the common ancestor of Dikarya. On the other hand, the analysis highlights that recent duplications are relatively rare. Likely, most duplications are rapidly destroyed by the action of RIP, as indicated by the high AT content of the pseudogenized alleles. For example, there is an unlinked, heavily mutated duplication of the gene *nwd6* in some strains, called *nwdp-2* (Figure 1). The gene *nwd6* itself can be pseudogenized too depending on the strain, suggesting that the duplication event that created *nwdp-2* triggered RIP for both copies.

Annotation of the N-terminal effector domains in both the NACHT and NB-ARC clades revealed that in the majority of cases the effector domains are related to confirmed cell-death inducing domains (Dyrka et al. 2020). Twenty of the 54 gene models have a predicted-lipase domain previously found to be involved in RCD execution, either the sesB domain (Daskalov et al. 2016) or the Patatin-like domain (Heller et al. 2018). Pore-forming domains related to the HeLo-like superfamily were also frequent and found in 15 gene models (Figure 1). When including the HET domain, the phosphorylase PNP-UDP domain, and the prion-forming domains (which in turn activate HeLo-like cell-death inducing domains) a total of 46 of the 54 studied gene models are associated to a domain with a confirmed or predicted cell-death inducing function (Paoletti and Clavé 2007; Daskalov et al. 2012; Rousset et al. 2023). This observation suggests that the majority of the NLRs in *P. anserina* function in the control of RCD.

As reported previously, there has been extensive shuffling of the N-terminal and C-terminal domains, relative to their ATPase domain (Figure 1 and **2**). In the case of NWD-related proteins, we can observe a variety of superstructure-forming domains typical of fungal NLRs: WD40, ANK, TPR, and HEAT repeats (Dyrka et al. 2014). By contrast, the NB-ARC genes are all associated with TPR repeats in *P. anserina* (Figure 2). The lack of phylogenetic resolution prevents the identification of clear transitions to different C-terminal repeat domains, except for the gene *Pa_1_15290* that has ANK repeats while being the sister group to the clade containing the HNWD genes, *nwd1*, *nwd3*, and *nwdp-2* (all with HIC WD40 repeats) (Figure 1). However, some transitions between different N-terminal domains can be confidently observed. For example, the *Pa_3_8560* gene has a PNP-UDP domain but is nested in a group of sesB-containing genes (Figure 1).

We used the program T-REKS (Jorda and Kajava 2009) to screen for repeats with HIC in the protein sequence of the superstructure-forming domains. T-REKS calculates a coefficient of similarity, *P_sim_*, representing an averaged Hamming distance of each individual repeat to a consensus (Jorda and Kajava 2009). Genes with a *P_sim_* of 0.7 or above were considered to have HIC, since lower *P_sim_* values are associated with higher false positive rates in the repeat detection (Jorda and Kajava 2009). Using this criterion, we determined that 15 of the 41 NWD-related genes have HIC, from which 11 have WD40 repeats, three have ANK repeats, and one has HEAT repeats (Figure 1 and **Table S2**). In the case of NB-ARC NLRs, seven out of 13 have HIC (Figure 2 and **Table S2**).

How general is the link between HIC and protein-length polymorphism based on repeat reshuffling? Dyrka et al. (2014) selected eight *P. anserina* NLRs with HIC (three NWD-related and five NB-ARC genes) and used PCR amplification in five strains from a Dutch population called the Wageningen Collection (van der Gaag et al. 1998), confirming that they all display length polymorphism. Expanding on this analysis, we extracted the ortholog of all 54 NLRs from the long-read genome assemblies of 13 strains from France and Wageningen (**Table S3**). Such long-read datasets are necessary to reconstruct HIC domains accurately (Ament-Velásquez et al. 2025). As expected, HIC and length polymorphism are perfectly correlated (Figure 1 and **2**). Thus, this class of repeats is at the same time internally conserved and hypervariable between strains, as concerted evolution (that is, HIC) represents both a cause and a consequence of repeat rearrangements. Correlation of HIC and repeat length polymorphism has also been observed previously in various types of protein repeats and phyla (Roche et al. 2003; Schüler and Bornberg-Bauer 2016).

In a phylogenetic context, our dataset illustrates the evolution of HIC amongst related genes with low internal conservation (LIC). For instance, the sister genes *Pa_7_10370* and *Pa_3_8560* have HIC in their ANK domain, while nested in a clade that has ANK repeats without HIC, or simply no size polymorphism (Figure 1). Moreover, although the WD40 repeats of *nwd5* and *nwd2* are closely related to those in the HNWD clade (Paoletti et al. 2007; Ament-Velásquez et al. 2025), the genes with HIC WD40 repeats were not resolved as monophyletic from the NACHT domain perspective. In addition, the gene *Pa_2_8180* has HIC ANK repeats but is related to an ANK gene with LIC (Figure 1). Despite the name, the WD40 repeats of *nwd4* (and related genes) are very divergent compared to those of other NWD genes, to the point of being unalignable, and display no HIC.

### Diversity, RIP-like mutations, and selection acting on NLR genes

To evaluate if the NLR genes evolve faster than other protein-coding genes, we first examined the fate of their orthologs in the genomes of six members of the *P. anserina* species complex (Ament-Velásquez et al. 2024). Although a single strain has been sequenced for some species, others have two to five strains available (**Table S3**), allowing limited inferences of intra-species diversity. For *P. anserina* we restricted the analyses to the 13 strains with high-quality genome assemblies, which are representative of the known diversity within the species (Ament-Velásquez et al. 2022) (**Table S3**). As a control, we randomly selected 100 protein-coding genes from the reference genome of *P. anserina* and manually curated their gene models as for the NLR genes, in order to make them comparable (herein, the “random” genes). We classified each gene as conserved (presumably fully functional), absent (deleted or absent from the genome), pseudogenized (based on stop codons, frame shifts, and TE insertions), polymorphic (present in some strains and absent or pseudogenized in others), or moved (translocated to another location in the genome) (Figure 3a and **Table S4**). The “moved” category is relatively easy to assess due to the very high collinearity and low divergence between the *Podospora* species (Ament-Velásquez et al. 2024). We found no cases where a gene had been moved in some strains but not others of the same species. Since the number of available strains varies across taxa, comparisons are more appropriate between gene classes rather than between species.

**Figure 3.**
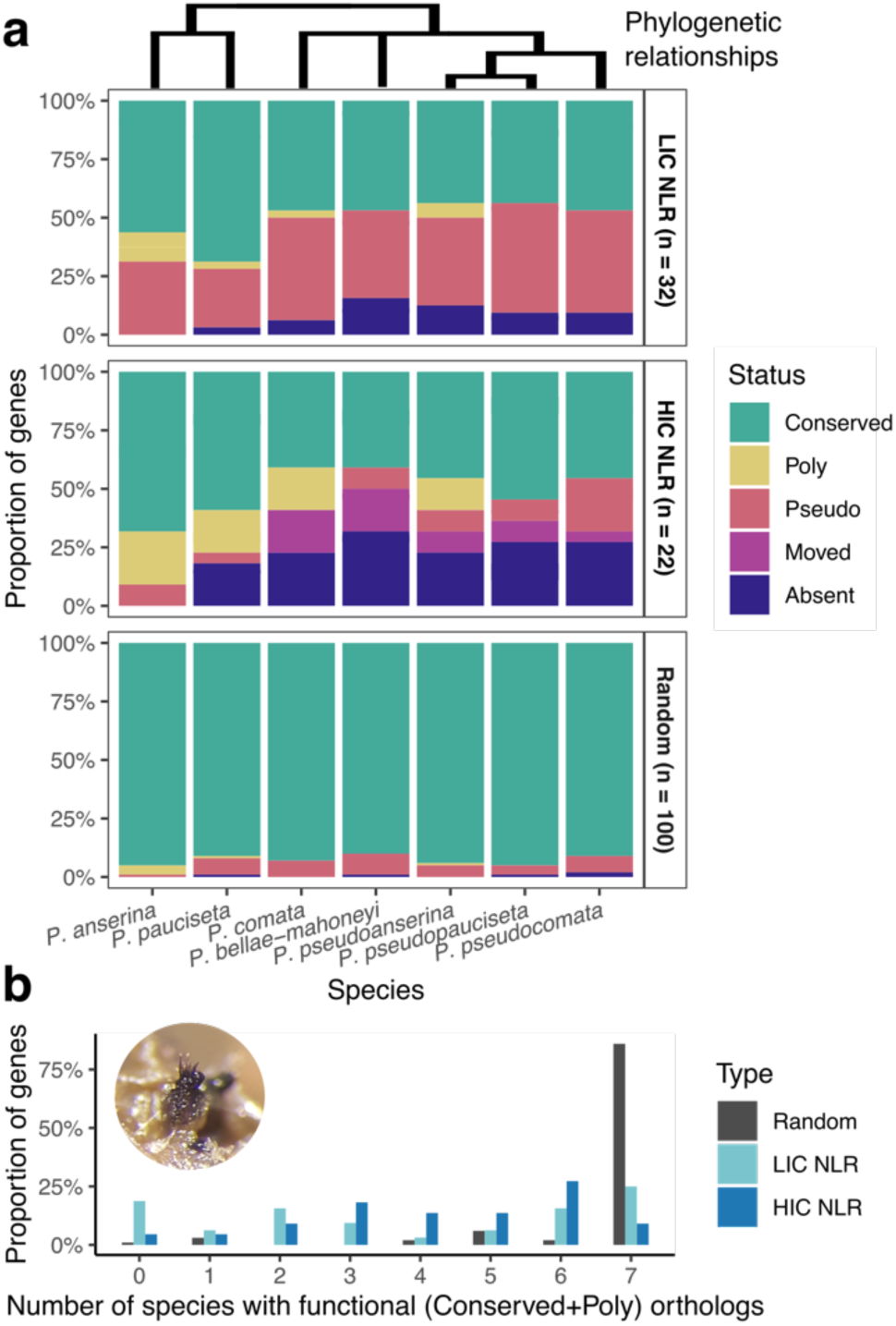
Conservation and distribution of NLRs and random genes in the *P. anserina* species complex. (**a**) Proportion of *P. anserina* genes at various degrees of conservation in every *Podospora* species for NLRs with low (HIC) or high internal conservation (HIC), and random genes. There is no Poly (present in some strains and absent or pseudogenized in others) category for species with a single strain available (*P. bellae-mahoneyi*, *P. pseudopauciseta*, and *P. pseudocomata*). Cladogram of phylogenetic relationships based on Ament-Velásquez et al. (2024). (**b**) Distribution of presumably functional orthologs across *Podospora* species. The inset image displays a close-up of a *P. anserina* fruiting body (Photo by S.L.A.-V.). **Figure 3 ALT TEXT.** Two graphs illustrating the degree of conservation of NLR genes in *Podospora* species labeled as **a** and **b**.

As expected, the random genes are highly conserved across the species complex (Figure 3a). By contrast, the NLR genes often fall in other categories, implying high gene turnover. Most random genes can be found in all seven species, but NLRs have a sparse distribution (Figure 3b). The HIC NLRs show particularly conspicuous presence/absence patterns between species, and unlike random genes and LIC NLRs, they are occasionally translocated to other regions in the genome (Figure 3a). Notably, HIC NLRs are significantly more associated with TEs based on the reference genome of *P. anserina* (Kruskal-Wallis rank sum [KW] test X^2^ = 36.414, df = 2, *p* = 1.238e-08, followed by post-hoc Dunn’s tests with Bonferroni correction [DB]) (Figure 4a), suggesting that TEs might be involved in the movement of these genes.

**Figure 4.**
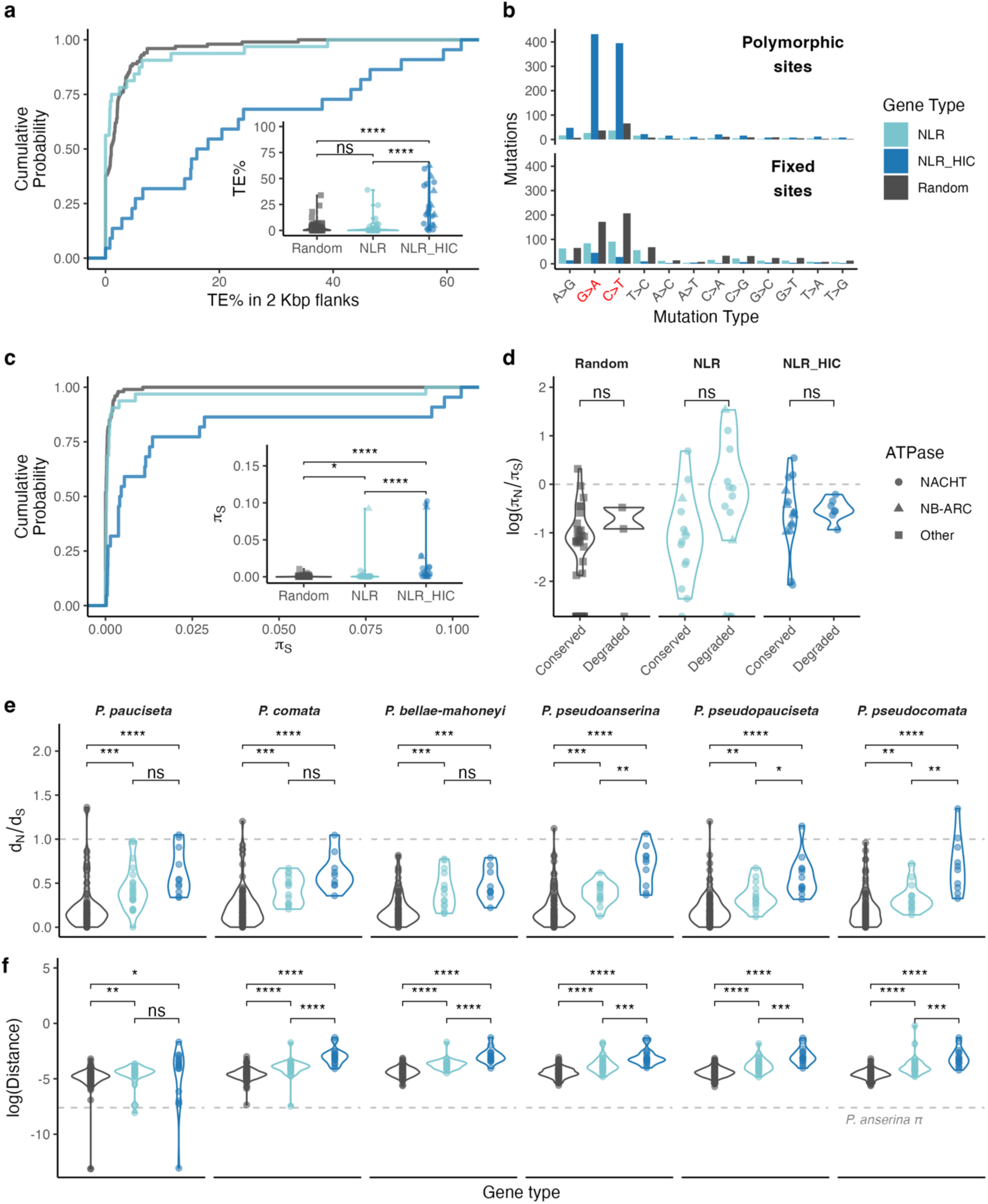
Contrasting evolutionary dynamics between random genes and NLRs. (**a**) Empirical cumulative distribution (ECDF) plot of the percentage of base pairs annotated as transposable elements (TE) in 2 kb-long flanks of each gene (4 kb in total), with distributions in a violin plot as inset. (**b**) Counts of mutations, polymorphic or fixed, within *P. anserina* gene types. The two transition types associated with RIP are highlighted in red. Mutation types are ordered such that transitions are first, followed by transversions. (**c**) ECDF plot of pairwise-nucleotide synonymous diversity (πS) for different gene classes, with distributions in a violin plot as inset. (**d**) Distribution of the ratio of non-synonymous (πN) to synonymous (πS) genetic diversity in *P. anserina* (log scale), making a distinction between genes that are presumably functional (conserved) or degraded in some form (pseudogenized or polymorphic). The dashed gray line marks πN = πS. In (**a**) to (**d**) different symbols are used to distinguish genes with NACHT and NB-ARC domains from other genes, as marked in (**b**). (**e**) Distribution of dN/dS values between *P. anserina* (strain S+) and the ortholog of the other species (type strains) for conserved genes. The dashed gray line marks dN = dS. (**f**) Phylogenetic distance measured as the path length (leaf to leaf) from the *P. anserina* gene to its ortholog in a given species for each gene phylogeny (all genes). The dashed gray line marks the average levels of genetic diversity (π) observed within *P. anserina*. Significance determined with Kruskal-Wallis rank sum tests followed by a *post hoc* Dunn’s test with Bonferroni correction (****: *p* <= 0.0001; ***: *p* <= 0.001; **: *p* <= 0.01; *: *p* <= 0.05; ns: not significantly different). **Figure 4 ALT TEXT.** Graphs comparing diversity and selection metrics of the three kinds of genes: random genes, LIC NLRs, and HIC NLRs. The subgraphs are labeled from **a** to **f**, and the comparisons include statistical values.

The close association of TEs and NLRs with HIC could imply a different input of RIP-induced mutations, which in *P. anserina* occur preferentially as C:G to T:A transitions mostly in the context of CpA:TpG and CpT:ApG dinucleotides (Hamann et al. 2000). We detected no significant difference in the median GC content between NLR types, although both have lower GC content than random genes (X^2^ = 65.171, *p* = 7.052e-15) (**Figure S1a**). We obtained similar results when excluding genes with signs of degradation (pseudogenized or polymorphic for the pseudogenized state) for both TE association (X^2^ = 30.183, *p* = 2.791e-07) and GC content (X^2^ = 44.257, *p* =2.452e-10).

To increase granularity, we focused on the GC content of polymorphic sites present in alignments of the coding regions. A sizable fraction of the diversity in NLRs is expected to be contained in the sensor domain (Van De Weyer et al. 2019). However, in the case of HIC NLRs, homology (identity-by-descent) cannot be assigned for the individual tandem repeats as the alleles have different repeat numbers presumably shuffled by recombination. Hence, we removed the HIC repeats except for the very first unit, since it tends to be slightly differentiated compared to the rest (and thus identifiable). We then inferred the ancestral state of all sites based on the sequence of the other *Podospora* species, whenever possible.

Despite removing most of their sensor domains, we found that HIC NLRs have many more polymorphic sites (997 mutations, n = 18 genes, or 55.4 mutations/gene) than LIC NLRs (137, n = 30, 4.6 mutations/gene) or random genes (163, n = 100, 1.6 mutations/gene). These sites are overwhelmingly dominated by C>T and G>A mutations, unlike the inverse transition directions T>C and A>G (Figure 4b and **Figure S1b**), strongly suggesting that they are methylation-or RIP-induced. Using a strict definition of a RIP-like site (C>T mutation proceeded by an A or T, or G>A mutation preceeded by an A or T) recovered a similar pattern (**Figure S1c**). Notably, this RIP dominance diminishes when counting fixed differences between *P. anserina* and the other species (Figure 4b and **Figure S1b**). To investigate this further, we performed a logistic regression using gene type (random, LIC NLRs, and HIC NLRs), mutation status of a site (polymorphic or fixed), and their interaction as predictors of whether a mutation was strictly RIP-like. The model revealed that, relative to random genes, LIC NLR genes exhibit marginally fewer RIP-like mutations among polymorphic sites (log-odds = –0.528, *p* = 0.041), while HIC NLRs showed a significant enrichment in RIP-like polymorphisms (log-odds = 0.942, *p* = 9.16e-08). In random genes, fixed mutations were significantly less likely to be RIP-like than polymorphic ones (log-odds = –0.447, *p* = 0.017). For HIC NLRs, the drop in RIP-like mutations from polymorphic to fixed was even stronger than in random genes (interaction log-odds = –0.755, p = 0.0059), while for LIC NLRs, this difference was not significant (interaction log-odds = 0.317, p = 0.29). Taken together, this pattern suggests that while RIP-like mutations arise frequently in HIC NLRs, they are rarely fixed, potentially due to purifying selection. The diversity and divergence of LIC NLRs, on the other hand, seem to be less affected by RIP than the genomic background.

As expected from the number of mutations described above, we found that the pairwise-nucleotide diversity at synonymous sites (π_S_), regardless of ancestral state, is higher for HIC NLRs compared to random genes (DB, *p* = 5.734e-11) but also to LIC NLRs (*p* = 0.0369) (Figure 4c). The same is true for non-synonymous sites (π_N_) (**Figure S2a**). We reasoned that degraded genes might have higher π_S_ due to relaxed selection. However, a Scheirer-Ray-Hare test (a non-parametric alternative to a two-way ANOVA) was only significant for gene type (H = 38.28, *p <* 0.0001). No significant effect of conservation status was observed (H = 0.524, *p =* 0.469), suggesting that whether a gene is conserved or degraded does not significantly influence synonymous site diversity in *P. anserina* (**Figure S2b**). This was also the case for the ratio of non-synonymous to synonymous diversity (π_N_/π_S_) (H = 32.981, *p <* 0.0001 for type; H = 1.429, *p =* 0.232 for conservation; H = 0.823, *p* = 0.663 for the interaction) (Figure 4d). Pairwise comparisons indicated that random genes have significantly lower π_S_/π_N_ values than LIC NLRs (DB, *p <* 0.0001) and HIC NLRs (*p <* 0.0001), but the two kinds of NLRs are not different (*p* = 0.2314). However, this observation should be taken with care, as the within-species π_N_/π_S_ metric is not a reliable indicator of the sign of selection (Kryazhimskiy and Plotkin 2008). Therefore, we turned to between-species comparisons.

We selected a representative sequence of *P. anserina* (namely the strain S+) and a representative of the other *Podospora* species, and estimated the pairwise rate of synonymous (d_S_) and non-synonymous substitutions (d_N_) per gene. A low ratio d_N_/d_S_ < 1 is generally regarded as evidence of strong purifying selection, whereas a high ratio d_N_/d_S_ > 1 indicates positive (“diversifying”) selection, and d_N_/d_S_ = 1 suggests neutral evolution (Jeffares et al. 2015). We found that both the gene type (H = 149.644, *p <<* 0.0001) and conservation status (H = 56.223, *p <<* 0.0001) influence the d_N_/d_S_ value of a gene. As expected, degraded genes have values closer to 1 (**Figure S3**). Hence, only conserved (presumably functional) genes in *P. anserina* were considered further (Figure 4e). Across all species comparisons with *P. anserina*, the random genes tend to have lower d_N_/d_S_ values compared to both kinds of NLRs (Figure 4e). There is also a weak tendency of HIC NLRs to have higher d_N_/d_S_ values than LIC NLRs (Figure 4e), although this could be an underestimation since most of the HIC sensor domain was excluded from the calculation. Overall, we rarely encountered values of d_N_/d_S_ > 1. However, the d_N_/d_S_ ratio calculated for a full gene is a relatively conservative metric, as most sites are under purifying selection, which dilutes the signal of a few positively selected sites (Jeffares et al. 2015). Hence, the lower d_N_/d_S_ in NLRs might suggest relaxed selection (although less than in their pseudogenized counterparts) or positive selection in some sites. The observed purging of RIP-like mutations in HIC NLR is in favor of the latter interpretation.

One expectation for both relaxed and positive selection is that NLRs should also have longer branches (i.e., evolve faster) than random genes in gene phylogenies. Indeed, our data confirmed that HIC NLRs are evolving faster than LIC NLRs, which in turn evolve faster than random genes (Figure 4f). However, comparing the phylogenetic distance distribution additionally revealed that some genes in *P. comata* and in particular *P. pauciseta* have suspiciously low levels of divergence compared to *P. anserina* (Figure 4f). As both balancing selection and introgression can result in lower divergence, next we assessed those possibilities.

### HIC NLRs display stronger signatures of balancing selection

In addition to the long-read assemblies, most of the *P. anserina* strains from the Wageningen collection (106 strains), were previously sequenced using Illumina technology and analyzed for population genomics statistics (Ament-Velásquez et al. 2022). That study revealed that the Wageningen collection and other sampled *P. anserina* strains from various locations exhibit extremely low genetic diversity across the genome (average species pairwise nucleotide diversity π = 0.000497). Furthermore, when the genome was divided into overlapping 10 kb windows, most genomic regions (96.25%) showed negative values of Tajima’s *D* (mean *D* of all windows = -1.62), an indication of pervasive purifying selection and/or a population expansion after a bottleneck (Tajima 1989). By contrast, several *het* genes were located within peaks of high diversity and positive Tajima’s *D* (Ament-Velásquez et al. 2022), consistent with balancing selection. We took advantage of these published estimates to calculate the average Tajima’s *D* values of the windows containing the NLR genes (Figure 1 and **2**).

We found that five NWD-related genes and four NB-ARC genes had positive Tajima’s *D* values, but the interpretation of this observation requires precaution. For example, the NWD-related gene with the highest positive *D* value is *nwd2* (Figure 1). However, this gene is located directly next to *het-s*, a well-studied allorecognition gene known to prevent the spread of deleterious plasmids (Debets et al. 2012). The *het-s* gene has two alleles, *het-s* and *het-S*, functionally defined by a single amino acid difference, position 33 (Deleu et al. 1993). A phylogenetic analysis including all *Podospora* species reveals polymorphism for the same amino acid in *P. comata* and *P. pauciseta*, and a topological split of the two alleles across the species complex, albeit with poor branch-support (**Figure S4**). In other words, the *het-s* phylogeny displays evidence of trans-species polymorphism. It has been reported that the *nwd2* variant linked to the reference *het-s* allele is interrupted by a *yeti* (aka. *repa*) LTR insertion (Deleu et al. 1990; Daskalov et al. 2012). We examined the genome assembly of all sequenced strains from the *P. anserina* Wageningen collection and confirmed that this TE insertion in *nwd2* is present only in the haplotypes with the *het-s* allele. The phylogeny of the *nwd2* gene mirrors that of *het-s*, although the *P. comata* alleles are recovered as monophyletic, suggesting some degree of recombination at *nwd2* within that species (**Figure S4**). Moreover, in some strains of other species, *nwd2* is fragmented or totally deleted. The presumed ancestral function of NWD2 was to activate HET-S in response to an unknown stimulus, but in *P. anserina* the mutation in position 33 resulted in a loss-of-function of the RCD-inducing activity of HET-S while at the same time allowing for self-activation through a prion state (Daskalov et al. 2015). Given the phylogenetic patterns of the other species, the allorecognition function of *het-s* likely predates the split of the species complex. As such, *nwd2* is at various degrees of pseudogenization in all or most lineages. Thus, the peak of high positive Tajima’s *D* probably reflects selection on *het-s* rather than on *nwd2* itself. A similar situation might apply to *Pa_1_11370* (Figure 2), which is linked to *PaPlp1* (part of the *het-z* locus).

Conversely, the known allorecognition genes *het-r* and *het-d* are located in windows with overall negative Tajima’s *D* values (Figure 1). However, their different alleles are defined by specific combinations of their HIC WD40 repeats (Espagne et al. 2002; Chevanne et al. 2009), which were filtered out of the SNP set used to calculate Tajima’s *D* (Ament-Velásquez et al. 2022). In fact, *het-r* has been shown to have balanced allele frequencies in the Wageningen population, while the interacting partner of *het-d*, called *het-c*, is multiallelic and under positive Tajima’s *D* peak itself (Ament-Velásquez et al. 2022; Bastiaans et al. 2014). Hence, these genes might still be under balancing selection despite their apparent negative Tajima’s *D*.

In addition to removing the HIC domain, the SNP window-based approach also filtered out regions with presence/absence patterns, which are more likely to contain NLRs (Figure 3a). Hence, as a complementary approach, we recalculated Tajima’s *D* from the gene alignments of the 13 *P. anserina* strains with high-quality assemblies (Figure 5a). As for the diversity metrics, these new estimates exclude most of the HIC domain, except for the very first repeat. Yet, we still found that the HIC NLRs have higher Tajima’s *D* values than random genes and LIC NLRs (KW test X^2^ = 17.258, df = 2, *p* = 0.000179; DB of Random vs. HIC NLRs *p =* 0.0001, LIC NLRs vs. HIC NLRs *p =* 0.0007, and LIC NLRs vs. Random *p* = 1) (Figure 5a). These relationships remain significant even if the entire HIC domain is removed all the way to the stop codon (X^2^ = 11.286, df = 2, *p* = 0.003542; DB of Random vs. HIC NLRs *p =* 0.0018, LIC NLRs vs. HIC NLRs *p =* 0.0072, and LIC NLRs vs. Random *p* = 1). Nonetheless, this approach has an additional limitation: it is likely an underestimation of the difference between NLRs and random genes, since 30 out of 100 random genes, but only one NLR, have no genetic diversity (π = 0) in our alignments. Tajima’s *D* cannot be estimated from zero diversity, but presumably a larger sample would result in negative values. Notably, while the Tajima’s *D* value of *het-d* remains negative (*D* = -1.789), it became slightly positive for *het-r* (*D* = 0.179) when including the first HIC repeat.

**Figure 5.**
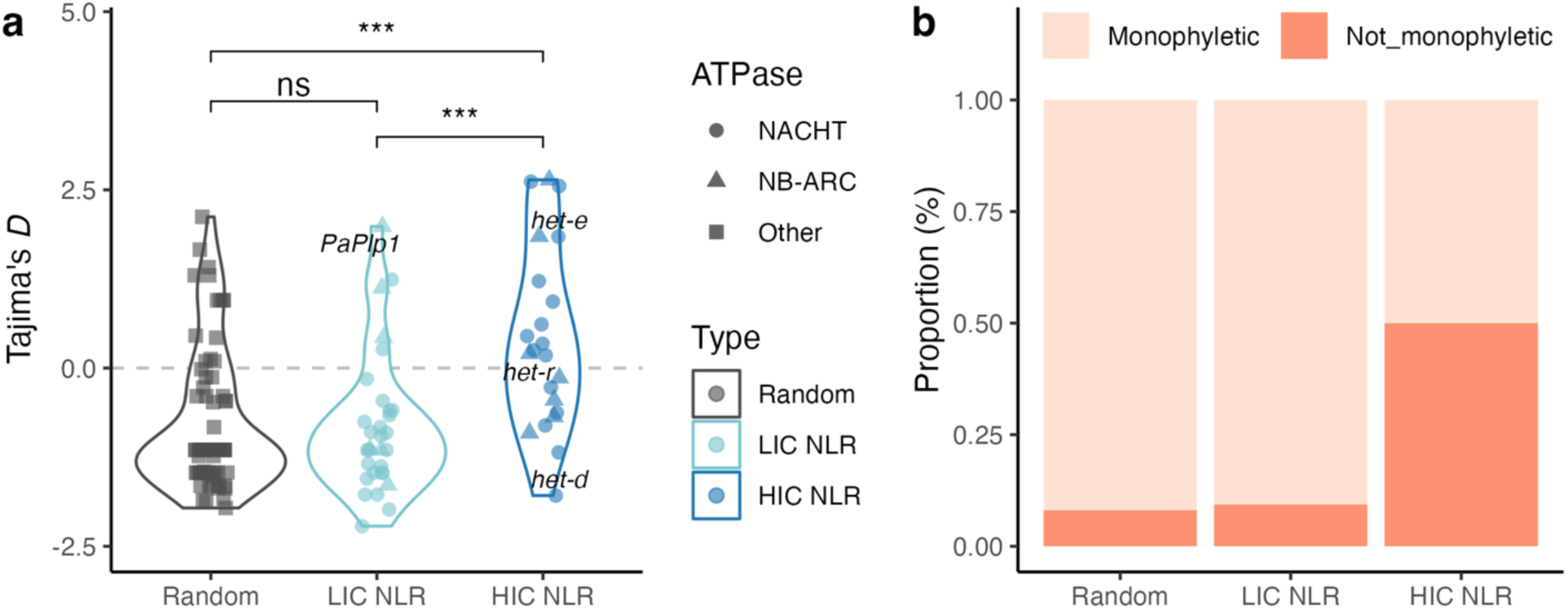
Balancing selection signals in random and NLR genes. (**a**) Distribution of the Tajima’s *D* statistic in the *P. anserina* strains with high-quality genomes for random genes, LIC NLRs, and HIC NLRs. Different symbols are used to distinguish genes with NACHT and NB-ARC domains (ATPase) from other genes. NLRs with an allorecognition function (*het* genes) are highlighted. Significance determined with Kruskal-Wallis rank sum tests followed by a *post hoc* Dunn’s test with Bonferroni correction (***: *p* <= 0.001; ns: not significantly different). (**b**) Proportion of genes where *P. anserina* is monophyletic in a gene phylogeny including all *Podospora* species. **Figure 5 ALT TEXT.** Two charts showing balancing selection in *Podospora anserina*. Panel a compares the Tajima’s D statistic across random genes, LIC NLRs, and HIC NLRs; panel b shows proportions of monophyly among these categories.

To evaluate possible long-term balancing selection, we produced phylogenies of all genes and asked how often *P. anserina* is non-monophyletic, as an indication of trans-species polymorphism. This analysis takes advantage of the fact that the *Podospora* species are very well defined as monophyletic clades separated by relatively long branches (Ament-Velásquez et al. 2024). We found that half of the NLRs with HIC are not monophyletic, while this proportion is much smaller for the other gene types (Figure 5b) (post hoc Fisher’s Exact Test after Bonferroni correction of HIC NLRs vs. random genes, *p* = 0.000052; HIC NLRs vs. LIC NLRs, *p* = 0.00398; LIC NLRs vs. random genes, *p* = 1).

Overall, there is more evidence of balancing selection driving the evolution of HIC NLRs than for LIC NLRs and random genes. However, introgression can be lurking behind some of these metrics.

### Patterns of genetic diversity and interspecies divergence support cases of both introgression and balancing selection on NLR genes

In order to explore the possibility of recent introgression, we estimated divergence (based on SNPs) between all 106 *P. anserina* strains sequenced with Illumina technology and the available strains from the other six *Podospora* species. We focused on chromosome 4 as a proxy of genome-wide variation to reduce computational time. We divided the chromosome in 2 Kb non-overlaping windows and calculated the pairwise-nucleotide differences to determine the distribution of divergence (**Figure S5a**). Windows with more than half their length missing due to unreliable SNP-calling were discarded. This analysis revealed that the average window divergence values between all 106 *P. anserina* strains and the other species are very homogeneous and well above 0.01 differences per bp (**Figure S5b**). However, there is an excess of low-divergence windows in comparisons between *P. anserina* and *P. pauciseta*, and to a lesser degree between *P. anserina* and some *P. comata* strains (**Figure S5a**). Therefore, we extended the analysis to the whole genome just for *P. pauciseta* and *P. comata*.

Divergence is strongly influenced, amongst other things, by the recombination and mutation landscapes (Noor and Bennett 2009; Ravinet et al. 2017). Both are unknown for any *Podospora* species. Still, as these are very closely related species with highly collinear genomes, we can expect their genomic landscape to be relatively conserved. Indeed, we found a strong correlation between the window divergence of *P. anserina* vs. *P. pauciseta* and the corresponding divergence of *P. anserina* vs. *P. comata* (e.g., n = 14814 windows, *r* = 0.596, Pearson’s correlation *p* < 2.2e-16 for comparisons with strain PaYp vs. either CBS237.71m or PcTdp) (**Figure S6**). In contrast, most regions with extremely low divergence in one species-pair are well differentiated in the other species-pair. Moreover, these low-divergence regions are located in different chromosomes depending on the strains compared (**Figure S6**). Taken together, these observations lead us to conclude that regions with divergence equal or below *P. anserina*’s intra-species diversity constitute recent introgression events. Defined as such, introgression is exceedingly rare: 2.07% - 1.49% of all the windows in comparisons with *P. pauciseta*, and 0.223% - 0.007% in comparisons with *P. comata*, depending on the strain.

These values are, however, underestimations, as this dataset was filtered such that every window is present in all strains (i.e., no missing data allowed). To get a more detailed view of divergence along the genome, we calculated 10 kb overlapping windows with steps of 1 kb (Ament-Velásquez et al. 2022), but we only removed windows that were composed of missing data in a given subset of strains: either *P. anserina*+*P. pauciseta* or *P. anserina*+*P. comata*. This scheme allows to detect windows that are present in one species pair, but absent in the other. Below we describe several cases of NLRs potentially involved in introgression and/or balancing selection.

The left arm of chromosome 4 is a particularly interesting case (Figure 6a). Most chromosomes have a relatively fast linkage disequilibrium decay when analysing the *P. anserina* Wageningen collection, with linkage disappearing completely well before the 10 kb distance in most cases (Ament-Velásquez et al. 2022). The exception is chromosome 4, whose decay can extend to around 40 kb (Ament-Velásquez et al. 2022). Notably, the divergence between the *P. anserina* and *P. pauciseta* strains collapses over more than 200 kb at the start of the left chromosomal arm, but not between *P. anserina* and *P. comata* (Figure 6a). The start of this region is in turn characterized by different divergence levels depending on the *P. anserina* strain. In other words, a large section of chromosome 4 was introgressed between *P. anserina* and *P. pauciseta*, most of it fixed in all the strains sampled (creating high linkage disequilibrium), but a section at the edge remains polymorphic, resulting in peaks of diversity and positive Tajima’s *D* (Figure 6a).

**Figure 6.**
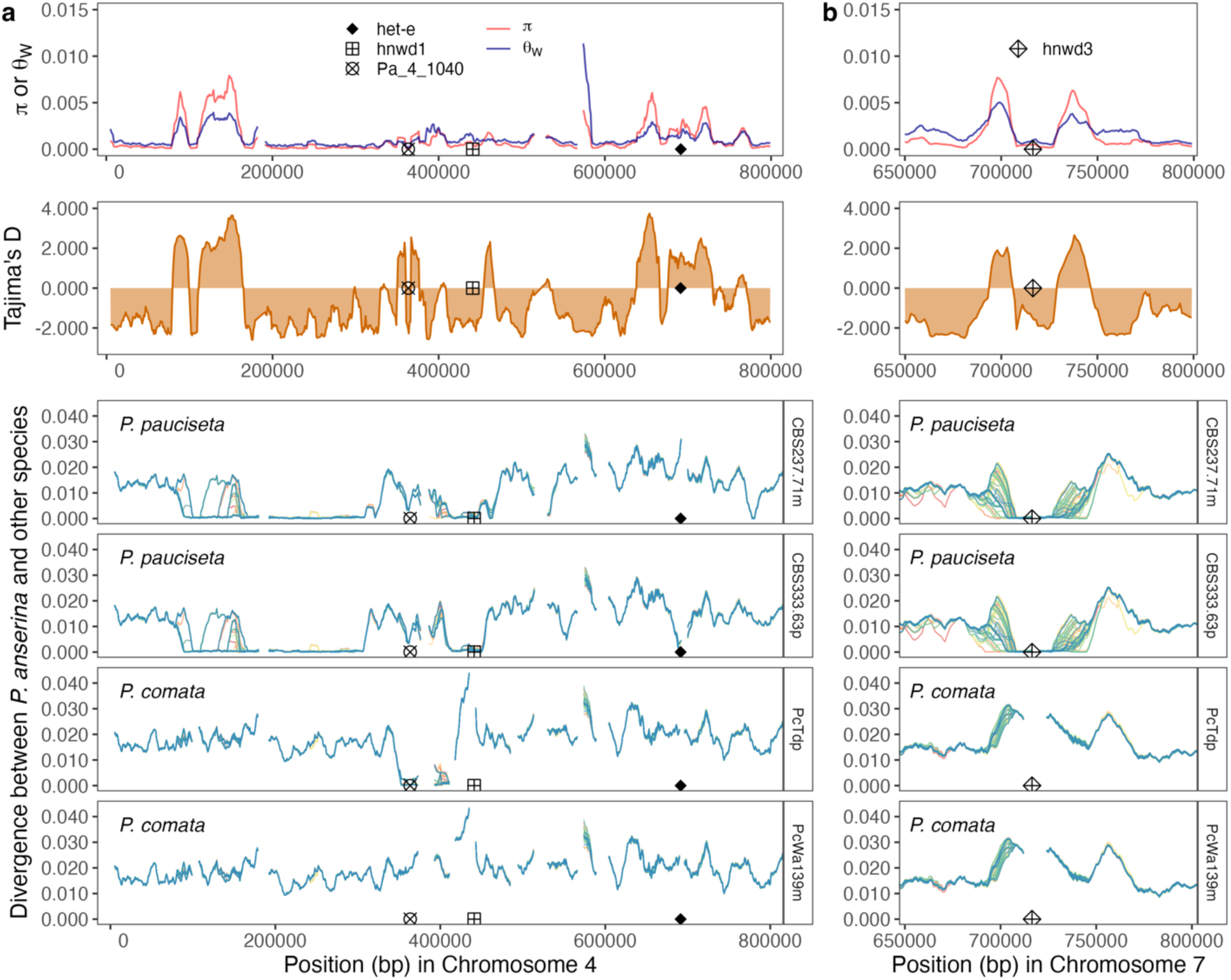
Population diversity and sequence divergence of regions containing NLRs in chromosome 4 and 7. The top panels correspond to the two metrics of genetic diversity π and Watterson’s θ as well as Tajima’s *D* in 10 kb windows (steps of 1 kb), as estimated in Ament-Velásquez et al. (2022) for the Wageningen population of *P. anserina*. The lower panels compare the pairwise-divergence between all 106 *P. anserina* strains vs. two different strains of either *P. pauciseta* and *P. comata*. Each *P. anserina* strain line was assigned a different color (gaps correspond to missing data). In (**a**) the plot corresponds to the start of chromosome 4, while (**b**) illustrates a close-up to the *hnwd3* region in chromosome 7. **Figure 6 ALT TEXT.** Multipanel figure showing different genome scans in selected regions of chromosome 4 to the left and chromosome 7 to the right.

Downstream of that region, several complicated scenarios appear for three NLRs: *hnwd1*, *Pa_4_1040*, and *het-e*. For these genes, divergence is narrowly reduced around them depending on the strain. In particular, the allorecognition gene *het-e* and its relative *hnwd1* seem to have been introgressed in some (CBS333.63p and CBS451.62p) or all *P. pauciseta* strains, respectively. Similarly, *Pa_4_1040* has been introgressed in three out of five *P. comata* strains. The same gene also has a relatively reduced divergence in the *P. pauciseta* strains (Figure 6a).

Another illustrative example comes from *hnwd3*, also closely related to *het-e*. This gene is flanked by peaks of diversity and Tajima’s *D*, but it has negative *D* values itself (Figure 6b). Importantly, *hnwd3* is exclusively present in *P. anserina* and *P. pauciseta*, and absent in the other five species. Phylogenomic analyses suggest that *P. anserina* and *P. pauciseta* are sister species (Ament-Velásquez et al. 2024). However, rather than common ancestry, this seems a case of introgression too: the *hnwd3* gene is nearly identical between the two species. Likely, the *hnwd3* gene was introgressed from *P. pauciseta* and underwent a selective sweep in *P. anserina*. The linked introgressed variation surrounding it has recombined with the original ancestral haplotype, creating a mosaic of ancestries of intermediate frequencies (hence the flanking Tajima’s *D* peaks), a pattern sometimes described as “volcano-shaped” (Moest et al. 2020; Setter et al. 2020). Chromosome 5 contains similarly complex regions sprinkled with introgression that involve NLRs, including *nwd5* and *PaPnt1* (**Figure S7**).

Given these examples, what is the expectation for a case of balancing selection without introgression? Consider *het-b*, a well-characterized (non-NLR) allorecognition gene with two divergent allelic classes (*het-B1* and *het-B2*) that are mutually monophyletic and with clear trans-species polymorphism involving *P. anserina, P. pauciseta, P. comata*, *and P. pseudoanserina* (Clavé et al. 2024). As expected, *het-b* nicely locates below a narrow peak of genetic diversity and Tajima’s *D* (**Figure S8a**). For such case, the divergence of the *P. anserina* strains compared to the interspecific *het-B1* strains (*P. pauciseta* CBS237.71-and *P. comata* Td+) and *het-B2* strains (*P. pauciseta* CBS333.63+ and *P. comata* Wa139-) varies depending on their allele, but it never goes below 0.01 nucleotide differences (**Figure S8a**). This observation, as well as the case of *het-s* discussed above (**Figure S4**), suggest that *bona-fide* allorecognition genes under long-term balancing selection are not expected to have extremely low divergence between species even within the same allelic class. Like these two allorecognition genes, there are NLRs with “clean” cases arguing for balancing selection. For example, *Pa_3_8170* has two marked allelic classes with highly divergent PNP-UDP terminal domains and is located under a narrow positive Tajima’s *D* peak with no indication of introgression (**Figure S8b**). The nearby NLR *Pa_3_8560* is under a similar Tajima’s *D* peak without signs of introgression (**Figure S8b**).

Lastly, some cases probably involved both introgression and balancing selection. A notorious example is the allorecognition locus *het-z*, which contains the NLR *PaPlp1* (Heller et al. 2018). As expected, the *het-z* locus is highly diverse and under a positive Tajima’s *D* peak (**Figure S9**). In our sample, the two *het-z* alleles are present in only *P. anserina* and *P. comata*, but the phylogenetic relationships of the alleles in the other species shows that the bi-allelic condition predates the split of the species complex (**Figure S9**), implying long-term balancing selection.

However, one of the two alleles (*het-Z2*) in *P. comata* has an extremely low divergence compared to the *P. anserina* allele, with the low divergence extending downstream (**Figure S9**). The alternative allele (*het-Z1*) is, by contrast, well differentiated and in line with the typical divergence between species. We interpret this pattern as evidence of introgression of a single allele for a locus already under balancing selection.

### Introgressed regions are enriched with NLRs

Given these numerous examples and the small proportion of areas involved in introgression, we wondered if NLRs were preferentially introgressed. We classified all windows into those that are involved in introgression with either *P. comata* or *P. pauciseta* (average divergence <= *P. anserina*’s π), windows discarded as missing data in both species, or non-introgressed windows. From this classification, it is evident that introgression is most common in chromosomes 4, 5, and 6 (Figure 7). There are 11655 protein-coding genes in our annotation, from which 571 are located in missing-data windows. From the 11084 surviving genes, 756 genes fall within the introgressed regions, including 13 NLRs. Due to their presence-absence patterns, 11/54 NLRs are located in the missing-data windows. Hence, 13/43 NLRs outside of missing-data windows have been introgressed. A Fisher’s Exact Test showed a significant difference in the proportion of NLR genes between introgressed and non-introgressed regions (odds ratio = 6.00, *p* = 3.359e-06).

**Figure 7.**
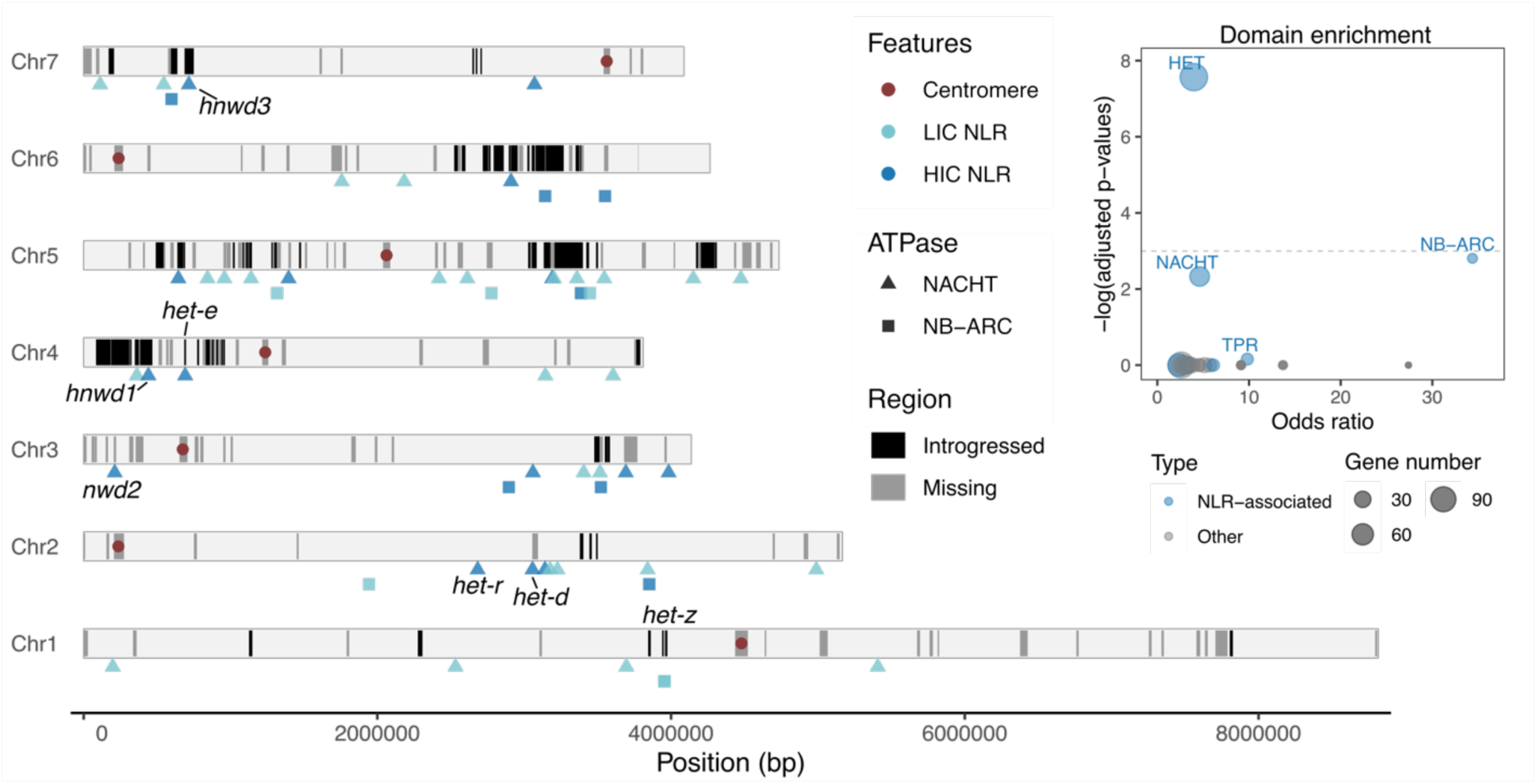
Chromosomal distribution of *P. anserina* NLRs in relation to introgressed regions from *P. pauciseta* or *P. comata*. The reference genome was divided in 10 Kb-long overlapping windows (1 kb steps), which were classified as either introgressed (black) or regions with more than half missing data (gray). The position of centromeres and the 54 NLRs are indicated, distinguishing between the two ATPase domains (NACHT and NB-ARC) and between standard NLRs and those with high internal conservation in their repeat domains (NLR HIC). Some of the genes discussed in the text are highlighted. In the upper right corner, an enrichment analysis based on PFAM annotations of genes with NLR-associated domains is shown. Domains that were not significantly enriched before multiple testing correction were omitted for clarity. The gray dotted line marks the significance threshold after Benjamini-Hochberg correction. **Figure 7 ALT TEXT.** Diagram of the seven *Podospora* chromosomes highlighting introgressed regions and the position of NLR genes. An insert to the right contains an enrichment analysis of domains in the introgressed regions.

As a complementary approach agnostic to our NLR dataset, we annotated the proteome using InterProScan, and we classified different NLR-associated domains based on their PFAM codes following Dyrka et al. (2014). An enrichment analysis as above revealed that out of all the identified domains in the proteome, the HET domain is significantly enriched in the introgressed regions (Figure 7 and **Figure S10**). After multiple-testing correction, the NB-ARC and NACHT domains are nearly significantly enriched (*p* = 0.061 and *p* = 0.097) (Figure 7).

### There is repeat exchange between HIC NLRs with TPR repeats

Recently, we characterized the relationships between the HIC WD40 repeats in *P. anserina* using a similar dataset (Ament-Velásquez et al. 2025). We found that some WD40 repeats of the *het-e* and *nwd6* genes are identical despite extensive protein differentiation in other domains, consistent with inter-gene repeat exchange. In a similar spirit, here we extracted the HIC TPR repeats of all NB-ARC NLRs from the 13 *P. anserina* strains with long-read data, amassing a total of 1024 repeats. We then performed a Principal Component Analysis (PCA) on their nucleotide sequences. While the repeats of most genes are highly differentiated from the others, those of *PaPnt1* and *Pa_7_3550* are largely equivalent in variation space (Figure 8 and **Figure S11**), with one shared identical repeat unit that occurs six times in *PaPnt1* and 13 times in *Pa_7_3550*. As these genes are otherwise quite divergent from each other in other domains (Figure 2), we interpret these observations as further evidence of repeat exchange between HIC NLRs. The three genes with ANK repeats are well differentiated and non-overlapping (**Figure S12**), but we expect interparalog exchange to occur in all repeat types.

**Figure 8.**
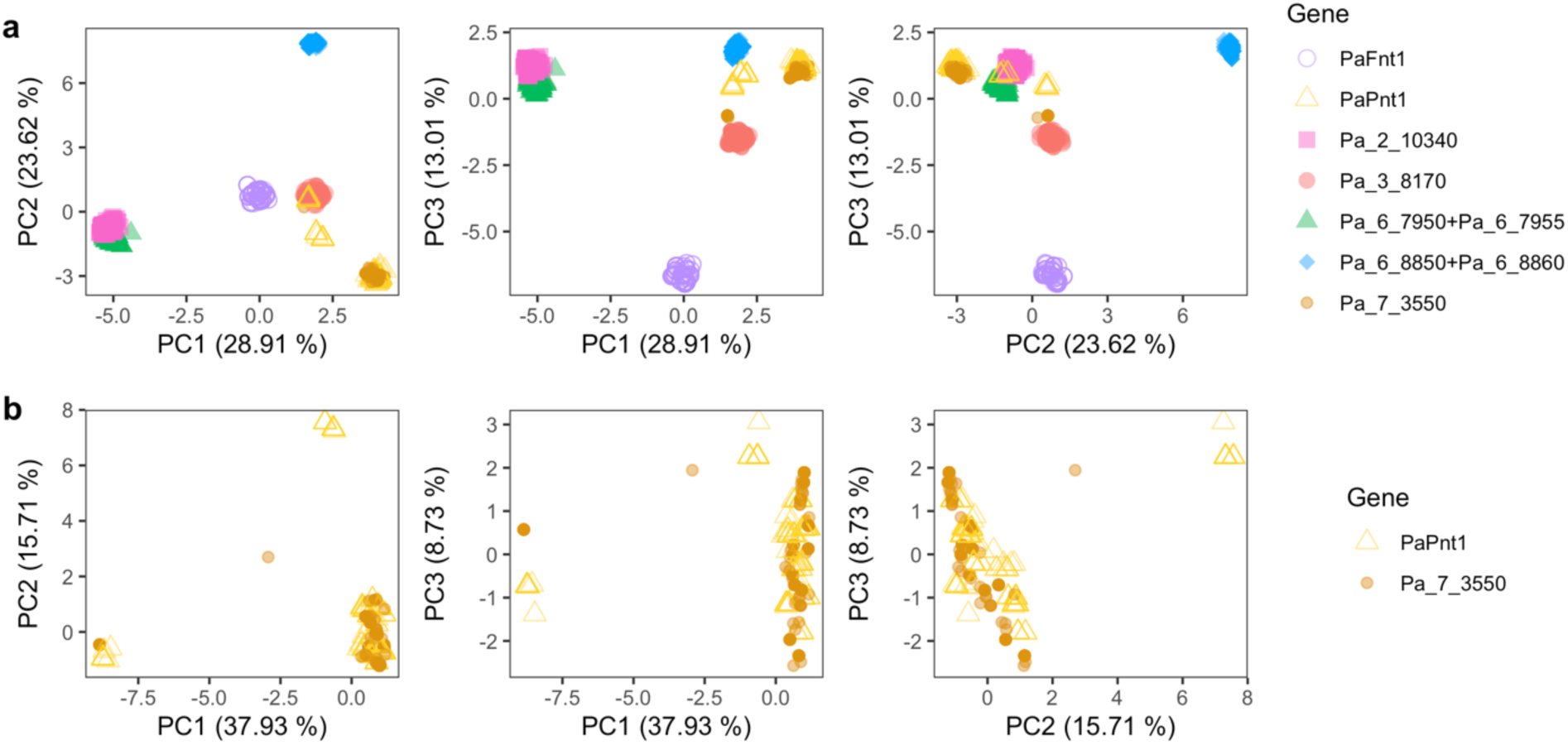
Principal component analysis of the TPR repeats with HIC. Considering all NLRs with HIC TPR results in strong differentiation for most genes (**a**). An exception is *PaPnt1* and *Pa_7_3550* which are largely overlapping (**b**). The genes *Pa_2_10340* and *Pa_6_7950+Pa_6_7955* are also overlapping but they are closely-related genes and hence expected to share similarities. **Figure 8 ALT TEXT.** Graphs of principal component analyses of TPR repeats with high internal conservation. The panel a contains all genes, while the panel b focuses on two specific genes.

### NLRs with HIC occur in all domains of life

Intrigued by the behavior of NLRs with HIC in *Podospora*, we decided to explore further the distribution of HIC NLRs across different taxonomic groups. Previous large-scale surveys found that HIC is rare in the proteins containing superstructure-forming repeats of Dikarya, Metazoa, and Viridiplantae (Dyrka et al. 2014). This observation is generally true across the Tree of Life for at least the proteins with WD40 repeats (Hu et al. 2017). However, there are indications that HIC is more common in NLRs for Dikarya, Metazoa, Viridiplantae, Actinobacteria, and Cyanobacteria (Dyrka et al. 2014; Daskalov et al. 2020). Hence, we extended these analyses to Archaea and to additional bacterial and eukaryotic groups (Figure 9). We chose the bacterial groups under the expectation that defense genes containing ATPases might be particularly abundant in multicellular lineages (Kaur et al. 2020). We extracted proteins annotated with ANK, TPR, WD40, and LRR repeats from the SMART database and inferred HIC using T-REKS as above. After reducing redundancy in the dataset, we found that indeed HIC is relatively rare amongst proteins of the four repeat types, confirming that internal conservation is by no means a structurally required feature for super-structure forming repeat domains (Figure 9a and **Figure S13**). However, from the proteins that do have HIC, many are associated with ATPase domains, i.e., they have an NLR-like architecture (Figure 9b and **Figure S13**). Specifically, among proteins without an ATPase domain, 5.59% contained HIC repeats, whereas 22.04% of proteins with an ATPase domain did. This corresponds to an odds ratio of 4.78 (95% CI: 4.51–5.06, Fisher’s exact test *p* < 2.2e-16). In the most dramatic cases, more than 30% of all HIC proteins in Ascomycetes, Basidiomycetes, and Actinobacteria have an ATPase domain (**Figure S14**). Notably, different lineages have larger repertoires of specific ATPase and HIC domain combinations. For instance, NACHT + HIC ANK proteins seem to be more of a Dikarya feature, while in Archaea, HIC proteins with an ATPase domain most often contain WD40 repeats. Finally, we manually scanned the proteins with HIC repeats and ATPase domains, and found multiple examples of full NLR architectures, often associated with N-terminal domains involved in RCD (Figure 9c). We conclude that while hypervariable HIC NLRs appear particularly common in Dikarya, this type of diversity-enhancing mechanism apparently also occurs in NLRs in other branches including animals, plants, and bacteria. Of note is the fact that, even in plant NLRs with low internal conservation, repeat shuffling has apparently governed NLR-family diversification in ancestral stages (Noël et al. 1999).

**Figure 9.**
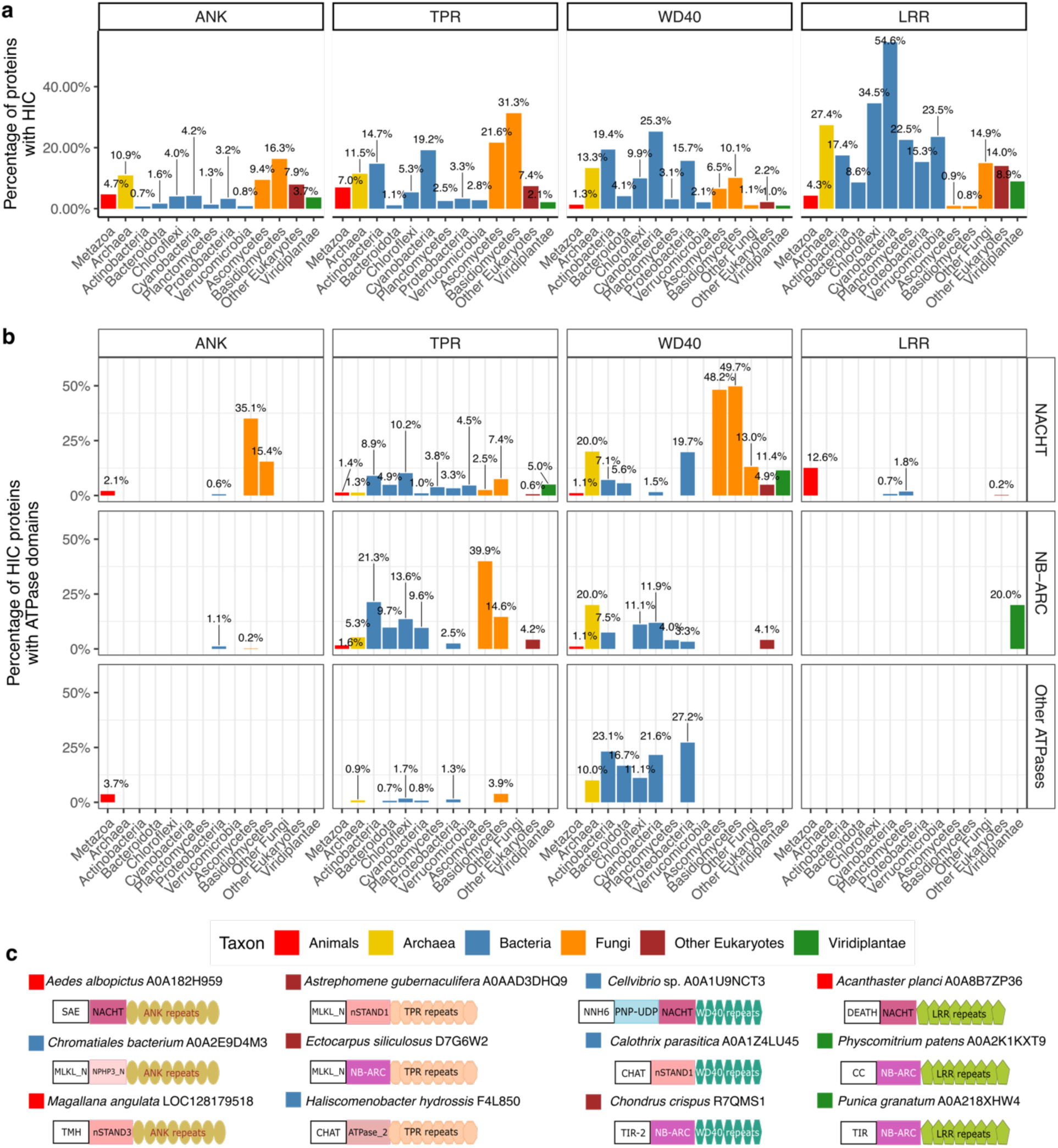
Distribution of proteins with HIC repeats across various taxonomic groups and repeat types. Proteins with HIC repeats are rare in general (**a**), but when present, they often contain an ATPase domain (**b**). In (**c**), manual curation of different selected proteins illustrate an NLR-like architecture for ATPase-containing HIC proteins. N-terminal domains no present in *Podospora* NLRs are represented with white boxes. Other ATPases: nSTAND1 (pfam: PF20702), nSTAND2 (PF20703), and nSTAND3 (PF20720) domains; DEATH: Death domain; CC: Coiled coil domain; CHAT: domain for Caspase HetF Associated with Tprs; MLKL_N: Mixed lineage kinase domain-like N-terminal domain; NNH6: NACHT N-terminal Helical domain 6; PNP-UDP: Phosphorylase superfamily; SAE: SAVED-associated Endonuclease; TIR-2/TIR: Toll-like receptors domain family; TMH: Transmembrane α-Helical Segment. Figure 9 **ALT TEXT.** Graphs showing distribution of high internal conservation across the Tree of Life. The panel a shows rarity across taxa, the panel b shows frequent ATPase association, and the panel c depicts curated architectures highlighting NLR-like structures and diverse N-terminal domains.

## Discussion

Filamentous fungi depend on anastomosis for growth and development, and hence rely heavily on allorecognition to preserve individuality, similarly to other modular organisms (Bastiaans et al. 2016; Aanen et al. 2008). In addition, fungal allorecognition provides protection against mycoviruses and a number of cytoplasmatic elements, acting as a form of immunity. However, it remains unclear if and how fungi detect additional pathogens or even potential mutualistic symbionts (Pawlowska 2024). The realization that several fungal allorecognition genes have an NLR-like domain architecture and trigger RCD cascades thus led to the hypothesis that the general function of fungal NLRs might be defense against pathogens, much like in plants, animals, and bacteria (Paoletti and Saupe 2009; Uehling et al. 2017; Daskalov 2023, 2025). In this study, we provided evidence that the bulk of *P. anserina* NLRs display evolutionary signatures consistent with an immunity function. Compared with other protein-coding genes, NLRs are more likely to have presence/absence patterns, have higher d_N_/d_S_ ratios, evolve faster, and are prone to introgression. NLRs with HIC repeats in particular seem to be more dynamic, as they have higher nucleotide diversity, collocate more with transposable elements, and readily exchange repeats between paralogs. Hence, this evolutionary perspective consolidates NLRs as prime candidates for genes that mediate fungal biotic interactions and offer concrete targets for molecular characterization.

The observed evolutionary dynamism of *P. anserina* NLRs was not an inevitable outcome, as these genes could have had entirely different functions with more selective constraints. For instance, the MalT protein in *Escherichia coli* has a NLR-like architecture but functions as a regulator of maltose metabolism that is unrelated to immune defense (Wu et al. 2023). The presence of N-terminal domains linked to RCD cascades and immune signaling in other taxa, however, suggest that most *P. anserina* NLRs control some form of programmed or regulated cell death. NLRs inducing RCD could play a role during morphogenesis or sexual development (Kufer and Sansonetti 2011; Bidard et al. 2024). Nonetheless, such scenario would not lead to fast change or require population polymorphism. Even among animals, where NLRs are firmly established as immune genes, evolutionary trajectories vary: in mammals, for instance, NLR repertoires are small and conserved relative to the diversity found in sponges, cnidarians, equinoderms, and some fish groups (Buckley and Dooley 2022). Overall, while some fungal NLRs may indeed carry out alternative functions, our results indicate that, on average, they experience selective pressures promoting divergence rather than strict conservation. This observation aligns well with a general immune function, understood in the broadest sense, as also including the regulation of potential symbiotic interactions in addition to pathogen defense.

Direct measurements of natural selection are extremely difficult to obtain and so we relied on the observed patterns of population diversity and divergence to infer the allele frequency dynamics. As a system, *P. anserina* offers an advantage to look for signatures of balancing selection: its high selfing rates increase the potency of background selection, removing neutral variation and making the sharp peaks of high polymorphism more obvious (Nordborg et al. 1996; Ament-Velásquez et al. 2022). Moreover, the clear differentiation between the *Podospora* species allowed us to distinguish trans-species polymorphism from recent introgression (at least to a degree). Both positive and balancing selection can facilitate introgression between species (Fijarczyk et al. 2016; Harrison and Larson 2014; Leducq et al. 2011). Accordingly, we detected several NLR genes embeded within areas that resemble selective sweeps after introgression., When treating them as a single functional class, we found that NLRs are enriched in introgressed regions, consistent with an adaptive interpretation. Additionally, we found higher average d_N_/d_S_ values in NLRs compared to random genes, although the estimates were close to 0. This result can be difficult to interpret on its own as higher d_N_/d_S_ could indicate both relaxed purifying selection or a mix of sites under strong purifying and diversifying selection. However, in combination with introgression patterns, occasional positive Tajima’s *D* estimates, and the apparent purging of RIP-like mutations during fixation, we argue that NLRs evolution in *Podospora* is more consistent with adaptation.

Since HIC is associated with allele size polymorphism at the population level in *Podospora* and at least one bacterium (Roche et al. 2003), it seems safe to assume that it constitutes a general variability-enhancing mechanism. Specifically, classic experiments in *P. anserina* demonstrated that loss-of-function mutations are overwhelmingly more common in the *het* HNWD genes than in their cognate ligands, driven by shuffling, gains, and losses of repeats (Labarère 1973; Chevanne et al. 2010). At the same time, different combinations of repeats can produce equivalent ligand affinities (Ament-Velásquez et al. 2025), potentially facilitating the exploration of the adaptive landscape. That said, there was no reason to expect higher nucleotide diversity in HIC NLRs outside their sensor domains. What exactly causes this increased diversity in *Podospora*? One source of mutations might be HIC itself. Recombination and repair mechanisms might have a role in the maintenance of HIC, creating additional *de novo* point mutations. Nonetheless, the most obvious source of variation seems to be RIP,which works by identifying highly similar (>80% identity) repetitive sequences that are more than ∼400 bp long (Galagan and Selker 2004). HIC NLRs with sufficiently long alleles, as well as those where there is interparalog repeat-exchange, might trigger RIP. The concerted evolution process that leads to HIC could subsequently remove mutations in the sensor domain, but adjacent domains might still get affected if RIP displays some leakyness (Van De Wouw et al. 2010). In addition, we found that NLRs with HIC are more associated to TEs, and hence could get occasionally duplicated via ectopic recombination of flanking TEs or even by direct mobilization, creating novel copies. Duplication leads to a strong RIP reaction, as observed in *nwd6* and its paralog *nwdp-2*. Finally, diversity might be higher simply due to positive or balancing selection overall, but that would imply that there is something particularly adaptive about the HIC architecture vs. other kinds of NLRs. We speculate that the combination of HIC and RIP is beneficial to create diversity faster, but it might lead to higher levels of autoimmune disorders. High variability implies that new variants might occasionally trigger unwanted RCD reactions, as observed in the NLR-induced hybrid necrosis of plants (Chae et al. 2014). Interestingly, the *het* NLRs in *Podospora* are known to cause self-incompatible offspring during outcrossing events, analogously to plant hybrid necrosis (Labarère 1973; Ament-Velásquez et al. 2022), but they can occasionally escape death via somatic mutation and deactivation of the HIC domain (Chevanne et al. 2010).

A large-scale analysis across Sordariales, the family of *Podospora*, found no evidence that RIP influences the number of NLRs per genome (Bonometti et al. 2025). This is paradoxical, given that NLRs have lineage-specific diversification events involving duplication, as discussed above. So, how are new NLR genes generated if RIP is there to destroy them? In principle, advantageous variants might survive in the long-term if selection is strong enough. However, there seems to be another route for NLR acquisition, as indicated by our results in *Podospora* – introgression. If an NLR evolved in a given species, its introduction into a new species via hybridization would not trigger RIP in the recipent, as long as its homologs are divergent enough. Horizontal gene transfer would have the same effect. Indeed, NLRs have been reported in transfer events between different fungal lineages (Qiu et al. 2016), and between fungi and bacteria (Ciach et al. 2024). Moreover, NLRs are often captured and mobilized by gigantic transposable elements known as *Starships*, which have also been implicated in horizontal gene transfers (Gluck-Thaler et al. 2022). In other words, obtaining diverged NLRs from other lineages can in part circumvent the limitations of having an active RIP system, which may otherwise limit evolutionary innovation (Galagan and Selker 2004). Ultimately, the final repertoire of NLRs for a fungus might be the result of their historical and present benefits, the interaction with genome defense mechanisms, vertical and horizontal lineage exchanges, and the genetic load imposed by these RCD-inducing fast-evolving loci.

The properties of HIC NLRs in *Podospora* are reminiscent of those observed in candidate immune receptors of brown algae. In these organisms, at least two main groups of receptors exhibit birth-death dynamics as observed here: one corresponding to NB-ARC+TPR NLRs, and another with a different domain architecture but including a superstructure-forming domain of LRRs (Zambounis et al. 2012). Intriguingly, in both gene classes the different repeats are encoded in individual exons, while the introns might contain additional repeats in the non-coding strand. Such a structure seems to facilitate exon reshuffling through alternative splicing, as well as by repeat exchange between paralogs (Zambounis et al. 2012; Teng et al. 2024). Similar to the case of *Podospora* HIC NLRs, it was hypothesized that the brown algae receptors might undergo somatic mutation, enhancing diversity beneficial for a potential immune function (Zambounis et al. 2012). Moreover, at least some of the brown algae receptors display a sort of weak HIC, where some repeats (including those on the non-coding strand) have high nucleotide similarity to each other. The working hypothesis in the brown algae field is that the repeat shuffling is a controlled mechanism, rather than a passive consequence of illegitimate recombination (Zambounis et al. 2012). Similarly, in the black truffle *Tuber melanosporum*, a distinct diversity-enhancing mechanism appears to operate on NLRs with ANK repeats that show extensive alternative splicing in tens of codon-sized mini exons (Iotti et al. 2012).

Nevertheless, HIC might be a more universal path to maintain polymorphism relative to these lineage-specific strategies. Zooming out to other taxonomical groups, large-scale surveys found that a significant proportion of NLRs in some groups of multicellular bacteria also show HIC (Daskalov et al. 2020). On the flipside, our survey found that HIC proteins often have an NLR architecture. That is, there seems to be an association between HIC and NLRs. Hence, it remains to be seen if the occurrence of HIC is controlled by common or multiple mechanisms across the Tree of Life and whether it offers adaptive benefits in other phyla in the context of self-nonself recognition.

## Methods

Most analyses were executed in Snakemake (Mölder et al. 2025) pipelines, with code available in the repository https://github.com/SLAment/MolEvoNLRs unless otherwise stated.

### Gene searches and annotation

Previous studies detected proteins related to the *hwnd* genes by doing BLASTp (Altschul et al. 1997) searches with a family member, in particular HET-E. This led to the discovery of the *nwd1*, *nwd2*, *nwd3*, *nwd4*, and *nwd5*, as well as the pseudogenes *nwdp-1* (here *nwd6*) and *nwdp-2*. To find more homologs, we performed similar searches on the NCBI website (https://blast.ncbi.nlm.nih.gov/Blast.cgi) while fixing the taxid to *P. anserina* (2587412). However, the BLASTp searches depend on good gene models that are often of bad quality for STAND proteins (Yuen et al. 2014). Hence, we performed tBLASTn searches with the NACHT domain protein sequence of HET-E (accession number XP_001903815.1, residues 294 to 738), as well as NWD4 (accession number CDP29667.1, residues 339 to 759), which has the most divergent NACHT domain relative to *het-e* from the known NWD genes. As reference, we used the Podan2 assembly from the S+ strain (Espagne et al. 2008), available on the Joint Genome Institute MycoCosm website (https://mycocosm.jgi.doe.gov/Podan2/Podan2.home.html). The tBLASTn searches were performed with the script query2haplotype.py v. 2.00 (https://github.com/SLAment/Genomics/blob/master/BLAST/query2haplotype.py) with options -- haplo --task tblastn -e 0.1, which fuses overlapping BLAST hits and defines individual haplotypes based on hits that are not farther than 10 kb from each other (controlled with the --vicinity option). We only kept resulting haplotypes that overlapped with at least one gene model from Podan2 and that had a recognizable core NACHT domain, including the ATP/GTPase P-loop (Walker A) motif, composed of the sequence GxxxxGK[ST], and the Mg2+-binding site (Walker B) motif (Koonin and Aravind 2000; Walker et al. 1982).

Similarly, we used the NB-ARC domain of PaPLP1 (accession number CDP23496.1, positions 371 to 672) as query to search for related NLRs with the same methodology as above. We confirmed that the NB-ARC domain contains the P-loop motif, composed of the sequence GxxGxGK[ST] (van der Biezen and Jones 1998). The gene *Pa_1_5560* was excluded from the results as it is likely pseudogenized in all species and there was no clear evidence to define the correct gene model. Additional genes were included if the NB-ARC domain (Pfam PF00931) was detected by InterProScan v. 5.62-94.0 (Jones et al. 2014) with default parameters.

We found multiple inconsistencies between the Podan2 gene models, more updated gene models of the S+ reference (Lelandais et al. 2022), an alternative annotation pipeline termed nice-3.00 (Ament-Velásquez et al. 2024), and the splicing sites suggested by RNAseq data of *P. anserina* (sample Psk2xS14-vsS mapped to the Podan2 assembly) or *P. comata* (samples PcTdp and PcWa131m mapped to the PcWa139m assembly) produced previously (Vogan et al. 2019). We gave priority to the RNAseq data patterns to choose or correct gene models amongst all the options when necessary. In the absence of RNAseq expression (<10x) in both *P. anserina* and *P. comata*, comparisons were made with the closest BLASTp hit in NCBI, which usually came from the genomes of *Apiosordaria backusii* strain CBS 540.89 and *Podospora setosa* strain CBS 892.96. Otherwise, the Podan2 gene model was taken as the correct one. Most of the inconsistencies were due to stop-codons, frame-shifts, or transposon insertions present in the S+ genome, which were often not there in homologs from other species detected by BLASTp in NCBI (last consulted on 2024-03-25). Hence, we marked the genes as potential pseudogenes. Occasionally, a given NLR was fragmented into two gene models in Podan2 due to such stop codons, frame-shifts, or transposons. In those cases the final gene model was re-named by fusing their IDs (e.g., *Pa_6_2410+Pa_6_2405*). After gene-model corrections, we kept a total of 41 NWD-related sequences (including the outgroup Pa_6_7270) and 13 NB-ARC NLRs for downstream analysis, although we detected at least 12 more pseudogenized remains of NWD relatives along the S+ genome that did not fulfill our NACHT minimum requirements.

Gene functional annotation was performed by inputting the corrected protein sequences to the NCBI Conserved Domains Database (Wang et al. 2023), as well as the hmmscan search engine (protein sequence vs profile-HMM database) at the HmmerWeb v. 2.41.2 (https://www.ebi.ac.uk/Tools/hmmer/search/hmmscan), with all “Protein Families” selected and default parameters (E-value = 0.01 and Hit = 0.03). Non-significant annotations were considered if they were consistent with known N-terminus domains in fungal STAND proteins (Dyrka et al. 2014). Overlapping annotations were merged for some Pfam categories with multiple variants (e.g., ANK repeats) following Dyrka et al. (2014) (**Table S1**). Classification of SesA and SesB sequences was further confirmed by comparing sequence alignments to the expected conserved motifs (Graziani et al. 2004).

To characterize the evolution of the NLRs at shallow evolutionary times, we retrieved the ortholog of each gene in all the *P. anserina* strains with long-read assemblies, as well as all available assemblies from the other known six species of the *P. anserina* species complex (**Table S3**). In this dataset, the genome of at least one representative of all species has been sequenced with long-reads (only one strain is available for *P. bellae-mahoneyi*, *P. pseudopauciseta*, and *P. pseudocomata*). We used the CDS of the reference Podan2 annotation (strain S+) as a query for the query2haplotype.py script to retrieve the nucleotide sequence of each gene, and confirmed orthology by examining the identity of the flanking genes. For every species, we classified the status of each gene as present, absent, pseudogenized, polymorphic, or moved. We inferred the movement (translocation) of genes based on comparisons of synteny of flanking genes, assuming that the chromosomal location that is most common amongst the seven species represents the ancestral state. We considered a gene pseudogenized if we observed frame-shifts, stop codons, or transposon insertions in the coding exons for most of the gene body except the last 50 amino acids of the C-terminus, under the assumption that changes in the position of the stop codon might still produce a fully functional protein. In the case of NLRs with HIC in the C-term domain, any frame-shifts or stop codons in the repeats themselves were ignored as the genes could be “resurrected” by repeat exchange. Loss of start codons was not considered a cause of pseudogenization if another methionine was located within 20 adjacent amino acids. Occasionally, some gene models would have a first exon extending farther away from the RNA-seq reads mapping area. In those cases, we looked for the next methionine downstream of the original start codon that was fully covered by RNA-seq reads and allowed stop-codons or frame-shifts to occur in the upstream exon region that lacked RNseq coverage (i.e. they were not marked as pseudogenes if mutations were present). As all strains with long-read assemblies of wildtype strains in our dataset have also been sequenced with Illumina Technology (Vogan et al. 2019, 2021; Ament-Velásquez et al. 2022; Hartmann et al. 2021), we verified all frame-shifts in SPAdes v. 3.12.0 (Bankevich et al. 2012) assemblies built with k-mers 21,33,55,77. Discordance between long-read and Illumina assemblies was typically located in runs of homopolymers, a known issue with these technologies (Delahaye and Nicolas 2021). In the case of samples that only had Illumina data available, the assembly often had tracks of Ns (missing data) within the repeat domain with HIC or was frequently fragmented into two or more contigs, making it impossible to assess mutations there. The alleles of the *nwd2* and *het-s* genes in the Wageningen Collection were extracted from SPades assemblies produced in the same way.

To contrast the dynamics of the NLR genes with other types of genes in the genome, we randomly selected 100 genes with CDS at least 1 kb-long (NWD-like family median size: 3.4 kb) from the Podan2 annotation and annotated them in the same way as above. For each gene, three sources of gene models were evaluated against the RNAseq data: the Podan2 and nice-3.00 annotations in *P. anserina*, as well as the equivalent annotation in the reference assembly of *P. comata* (strain Td+, assembly PODCO) (Silar et al. 2019). The gene model of Pa_5_340 was ambiguous but we confirmed that the N-terminus domain is truncated by comparing its AlphaFold model to close homologs (e.g., KAK0659194.1) and considered it a pseudogene. In the case of Pa_7_10370, the stop codon in the Podan2 and nice-3.0 models locates within the ANK domain, while other species have a stop codon downstream outside any HIC repeat. An earlier stop codon in *P. anserina* is also present in an ANK repeat but within a small intron that is not supported by RNAseq data. While this implies pseudogenization, the RNAseq expression is considerable (∼100x) and decays after the first stop codon. If the gene of a given strain was not fully assembled or the read mapping was not conclusive, it was considered missing data (e.g., Pa_7_11560 in Wa131-, Pa_3_4260 in Wa53-, and Pa_7_11300 in CBS333.63+).

The final gene models of all NLRs and random genes were included in an updated version of the “nice” annotation (Ament-Velásquez et al. 2024), termed nice-3.02 and available as a gff3 file in the GitHub repository.

### Phylogenetic analyses

Protein sequences were aligned with the online version of MAFFT 7 (Katoh and Standley 2013) with default parameters (BLOSUM62 scoring matrix, gap penalty of 1.53). We used IQ-TREE v. 2.2.3 (Nguyen et al. 2015) with automatic selection of substitution model (-m MFP) and 100 bootstrap pseudo-replicates (-b 100) to produce maximum likelihood phylogenies. Details about the specific alignments are given below.

The general phylogeny of the NWD-like genes in *P. anserina* was produced from the core region of the NACHT domain (Koonin and Aravind 2000), corresponding to the positions 294-629 in the HET-E protein sequence of the strain S+ (GenBank accession number CDP27944.1). We used the Pa_6_7270 gene as an outgroup. In general, we used the reference strain S+ sequence of a given *P. anserina* gene or protein with the exception of *nwd6* (pseudogenized in S+), for which we used the allele in Wa21-(assembly PaWa21m). The phylogeny of the core region of the NB-ARC domain (van der Biezen and Jones 1998) was done as above, with an alignment that corresponded to the positions 404-753 of PaPLP1 (CDP23496.1). In the absence of an outgroup we resorted to midpoint rooting in FigTree v. 1.4.4 (http://tree.bio.ed.ac.uk/software/figtree/).

### Protein repeat analysis

We used the online version of T-REKS (Jorda and Kajava 2009) with default parameters to identify repeats with HIC (last consulted in August 2024; https://bioinfo.crbm.cnrs.fr/index.php?route=tools&tool=3). T-REKS infers the boundaries of tandem repeats in a given protein and produces a sequence alignment of the repeats. Then it estimates repeat similarity by calculating the Hamming distance *D_i_* between each repeat and the overall repeat consensus. Namely, for an alignment of length *l* and *m* repeats, a similarity coefficient is defined as: 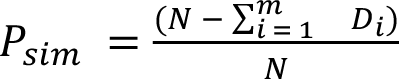, with *N* = *m* × *l* and 0 ≤ *P_sim_* ≤ 1 (Jorda and Kajava 2009). In a similar spirit to that of Hu et al. (2017), we use *P_sim_* = 0.70 as a threshold to consider repeats as internally conserved, which also happens to be the minimum *P_sim_* value of the online T-REKS tool. We considered a protein as LIC if T-REKS failed to detect repeats. As input we used the allele of the S+ strain, except for genes that were pseudogenized, absent, or had less than three repeats (*het-e*, *nwd5*, and *nwd6*), in which case we used the allele from the Wa21-strain.

Preliminary testing of T-REKs output showed inconsistencies in the inferred number and boundaries of the repeats. Thus, we extracted the alleles of all the *P. anserina* long-read assemblies and aligned them manually. We defined HIC WD40 repeats by following the structure of the WD40 canonical repeat (Hu et al. 2017). ANK, TPR, and HEAT repeats were defined by following the consensus of Al-Khodor et al. (2010), Marold et al. (2015) and Yoshimura & Hirano (2016), respectively.

In order to systematically extract individual HIC repeats in all strains, we implemented a simple strategy based on regular expressions (regex) of repeat types (Ament-Velásquez et al. 2025). Specifically, we used a regex string to retrieve repeats along each allele from individual protein sequences (available in the GitHub repository). If an allele had frame-shifts mutations, gaps were introduced to restore the coding frame and the codon was treated as a stop codon before applying the regex. As previous studies have shown that *het-r* and *het-e* functional alleles can be deactivated with as little as one amino acid INDEL (Chevanne et al. 2010), we only retained repeats that were 42 (WD40 repeats), 33 (ANK and HEAT) or 42 (TPR) amino acids long. However, manual alignment of the ANK repeats of Pa_2_8180 and Pa_7_10370 revealed repeat units with non-canonical lengths that appeared consistently in multiple strains. Therefore, we allowed for a range of 34 or 35 amino acids for Pa_7_10370 or 33 to 36 amino acids for Pa_2_8180 to define full repeats. Stop codons were allowed as long as they didn’t conflict with the regex. Potential cryptic repeats (not HIC) and occasional half repeats that appeared at a fixed position in nearly all strains (e.g., in Pa_2_8180) were discarded.

From the obtained HIC repeats collection we re-calculated *P_sim_* in python using the output of the gap_consensus(threshold=0) method of the AlignInfo object in the biopython library v. 1.85 (Cock et al. 2009). These new *P_sim_* correspond to those reported in Figure 1. The protein sequences of the repeats were aligned with ClustalW v. 2.1 (Larkin et al. 2007) to produce a sequence logo per gene, plotted with the R package ggseqlogo v. 0.2 (Wagih 2017). The nucleotide sequences of the TPR repeats were read into R with the ape package v. 5.7 (Paradis and Schliep 2019). We produced Principal Component Analyses with the package adegenet v. 2.1.10 and ade2 v. 1.7-22 (Jombart 2008).

### Population genomics analyses

To explore the patterns of diversity of the NLR genes at the population level, we used published window estimates (10 kb-long with steps of 1 kb in the S+ reference genome) of π, Waterson’s θ, and Tajima’s *D* in the Wageningen Collection (Ament-Velásquez et al. 2022). We averaged the Tajima’s *D* value of all the windows that overlap with the model of each gene to obtain a proxy for the gene-specific values. Since each window has a certain amount of missing data, we discarded windows that have less than 5000 bp of sites genotypified (Ament-Velásquez et al. 2022).

We used the CDS nucleotide alignment of all genes (random and NLRs) extracted from long-read assemblies to infer several gene properties in *P. anserina*. As the direct homology (identity-by-descent) between individual HIC repeats is unknown, we removed the HIC repeats, leaving only the very first repeat of each NLR as a diversity representative, which is often slightly more differentiated than the subsequent repeats. Tests excluding this repeat and the full C-term yield similar results (not shown).

We counted the proportion of each nucleotide across all sequences in the alignment to produce averaged GC proportion estimates. We used the python library EggLib v. 3.1.0 (Siol et al. 2022) to calculate the pairwise nucleotide diversity for all sites (π), non-synonymous sites (π_N_), synonymous sites (π_S_), and the ratio π_N_/π_S_ based on the counting method of (Nei and Gojobori 1986). After defining codons with egglib.tools.to_codons, we calculated the statistics by setting the parameters multiple_alleles=True, multiple_hits=True, max_missing=1, skipstop=False in the CodingDiversity call, effectively tolerating all missing data and treating stop codons as another amino acid.

We ran RepeatMasker v. 4.1.5 (http://www.repeatmasker.org/) with the library “PodoTE-1.00” (Vogan et al. 2021) to annotate repeats on the Podan2 reference (strain S+). We then calculated the proportion of base pairs annotated as TEs in 2 kb flanks of each focal gene. We determined the coordinates of 2 kb before the start codon and in 2 kb after the stop codon and intersected them with the TE annotation using BEDtools v. 2.31.1 (Quinlan and Hall 2010).

For each gene, we extracted the corresponding ortholog from the available assemblies of the other *Podospora* species and aligned them together with the *P. anserina* sequences. We extracted the variable sites and determined if they were polymorphic in *P. anserina*, or fixed and different from the other species. We chose biallelic sites and assigned the ancestral state based on the nucleotide present in the six other *Podospora* species. Triallelic sites or sites where at least one *Podospora* species disagreed were ignored to avoid ambiguities. We also recorded the nucleotide bases flanking said mutations. A transition C>T was considered RIP-like if an A or T was located downstream (ancestral CpA/T sites). Likewise, G>A was considered RIP-like if an A or T was upstream (reflecting GpA/T sites in the antisense strand). We treated RIP-like as a binary variable and applied a logistic regression with the gml() function in R to investigate whether the frequency of RIP-like mutations differed among gene categories and mutation statuses (fixed or polymorphic).

We created CDS alignments with a single representative of all *Podospora* species by selecting the type strain of each taxon (**Table S3**) whenever possible, otherwise a random strain was used. If the stop or start codons were different between species, alignments were trimmed to match the model in *P. anserina.* We used Q-TREE as above to produce phylogenies of all genes with at least three taxa. We used the tree.distance call of the biopython library to calculate the distance between the *P. anserina* sequence and each species. The same alignment was used to estimate d_N_/d_S_ between the *P. anserina* ortholog and each of the other six species based on the model-averaged method (MA) implemented in KaKs_Calculator2 v. 2.0.1 (Wang et al. 2010; Zhang et al. 2006). Two comparisons in genes introgressed between species pairs with zero synonymous sites (creating values of KaKs = 50) were discarded.

We estimated divergence along the genome between all published 106 *P. anserin*a strains and each strains from the other species with a modified version of the variant calling pipeline of Ament-Velásquez et al. (2022). Using the Podan2 assembly as reference, we mapped the paired-end Illumina reads of all strains with BWA v. 0.7.17 (Li and Durbin 2010), followed by PCR-duplicates marking with Picard v. 2.19.0 (https://broadinstitute.github.io/picard/). We used the HaplotypeCaller of the Genome Analysis Toolkit (GATK) v. 4.5.0.0 (Poplin et al. 2017) for joint variant calling, as recommended by the GATK Best Practices (Van der Auwera et al. 2013). We applied a hard-filtering scheme of QD < 2, FS > 60, MQ < 40, QUAL < 30, SOR > 3, ReadPosRankSum < -8. Only surviving SNP variants in the nuclear chromosomes were consider further (i.e., mitochondrial data was ignored). We extracted the depth of coverage across all sites for each sample using the package vcfR v. 0.1.16 (Knaus and Grünwald 2017) and discarded sites that were below the 25% quantile or above the 98.5% quantile depth distribution (Ament-Velásquez et al. 2022). All SNP variants overlapping with annotated TEs were also discarded.

Next we applied different filtering schemes depending on the analysis. First, we aimed at estimating general levels of introgression between the seven species. However, as this is computationally demanding, we focus only in chromosome 4 as a representative. We discarted all sites with missing data using VCFtools v. 0.1.16 (Danecek et al. 2011) with parameter --max-missing 1 to make the data comparable across taxa. We ran the R package PopGenome v. 2.6.1 (Pfeifer et al. 2014) to calculate pairwise-nucleotide divergence between pairs of strains (i.e., each *P. anserina* strain vs. each strain of other species) in non-overlapping windows of 2 kb. Windows with less than 1 kb of genotyped sites (variable and invariable) were discarded. To calculate introgression proportions, we re-ran the pipeline (including site filtering) for the whole genome of *P. anserina*, *P. pauciseta*, and *P. comata*. Finally, we re-ran the pipeline for the three species but defining 10 kb overlapping windows with steps of 1 kb, discarding windows with less than 5 kb of data (Ament-Velásquez et al. 2022). In this case, filtering was done by species pairs (*P. anserina* with either *P. pauciseta* or *P. comata*), such that all strains of each species-pair had the same windows. We further classified all the windows with an average divergence between either species-pair equal or smaller than *P. anserina*’s intra-species diversity (π = 0.00049) as introgressed. Windows that were discarded as missing data in both species comparisons were set as “missing-data regions”.

All statistical analyses were performed in R v. 4.4.2 with the packages tidyr v. 1.3.1 (Wickham et al. 2025b), dplyr v. 1.1.4 (Wickham et al. 2025a), rcompanion v. 2.5.0 (Mangiafico 2025), dunn.test v. 1.3.6 (Dinno 2024), and stats v. 4.4.2 (R Core Team 2021). Plotting was done with the packages ggplot2 v. 0.6.0 (Wickham 2016), ggpubr v. 0.6.0 (Kassambara 2023), cowplot v 1.1.3 (Wilke 2024). .

### Enrichment analyses of PFAM codes in introgressed regions

We performed an enrichment analysis to determine whether specific PFAM protein domains were significantly overrepresented in the introgressed regions compared to the rest of the proteome. As reference we used the protein-coding annotation in Ament-Velásquez et al. (2024) modified with our manual curation of the NLRs and random genes. We ran InterProScan v. 5.75-106.0 (Jones et al. 2014) to obtain PFAM annotations. If the CDS of a gene overlaped the introgressed regions, it was considered introgressed. Conversely, only genes whose CDS was fully contained within missing-data regions were classified as missing, ensuring the two datasets were mutually exclusive. For each PFAM code, we counted how many domain-containing genes overlap with the introgressed regions and how many did not, while excluding the genes in missing-data regions in both cases. We employed Fisher’s exact test for each PFAM contingency table. To account for multiple hypothesis testing, we adjusted the resulting *p*-values using the Benjamini-Hochberg procedure. PFAM codes with adjusted *p*-values below a significance threshold of 0.05 were considered significantly enriched in the introgressed regions. Notice that, while earlier versions of InterProScan classidied NACHT proteins as the PFAM PF05729, the latest version classifies most of them as PFAM PF24883 (NPHP3_N). Here, we re-classified them as NACHT (**Table S1**).

### Looking for HIC across the Tree of Life

Following Daskalov et al. (2020) we extracted proteins annotated for WD40, ANK, TPR, and LRR repeats from the SMART (Letunic et al. 2021) database (last consulted in Jun 2025) for various taxonomic groups. Redundancy of fasta files was reduced using the online version of DIAMOND-DeepClust v2.1.3.157 (Buchfink et al. 2023) (available at https://toolkit.tuebingen.mpg.de/tools/diamond_deepclust) with default parameters (minimum identity percentage and minimum coverage percentage of 80%). We then ran T-REKS with default parameters to infer repeats and calculate the *P_sim_* metric. For each sequence we discarded repeat units inferred to be smaller than 21 and bigger than 100 bp (repeat types are usually around 40 bp but T-REKS sometimes identifies two tandem repeats as a unit). From the resulting repeat inferences, we chose the one with the largest coverage in the protein as a representative. A protein was inferred to have HIC if the chosen representative repeat region had a *P_sim_* >= 0.70. We re-annotated the surviving proteins (with and without HIC) using InterProScan with default parameters and parsed the output to identify the presence of AAA+ ATPases (NACHT, NB-ARC, nSTAND1, nSTAND2, and nSTAND3). Proteins with more than one type of superstructure-forming repeats were discarded due to the ambiguity of assigning HIC automatically (0.51% of the proteins in the final dataset of 160477 proteins).

## Data availability

The source of all used genome assemblies is available in **Table S3**. A collection of Snakemake pipelines is available at https://github.com/SLAment/MolEvoNLRs.

## Acknowledgments

We thank Brendan Furneaux and Sergio Tusso for helpful discussions. This work was supported by the Swedish Research Council (grant 2022-00341) and the Stiftelsen Anna-Greta och Holger Crafoords fond (CR2023-0039) to S.L.A.-V. The computations were performed on resources provided by NAISS at Uppsala Multidisciplinary Center for Advanced Computational Science (UPPMAX) partially funded by the Swedish Research Council through grant agreements no. 2022-06725 and no. 2018-05973. We would also like to thank the PDC Center for High Performance Computing, KTH Royal Institute of Technology, Sweden, for providing access to the computing resources used in this research.

## Notes

https://github.com/SLAment/MolEvoNLRs

### Competing Interest Statement

The authors have declared no competing interest.

