## Supplementary Figures for "NOD-like receptor genes evolve under diversity-enhancing mechanisms in a fungal species complex"

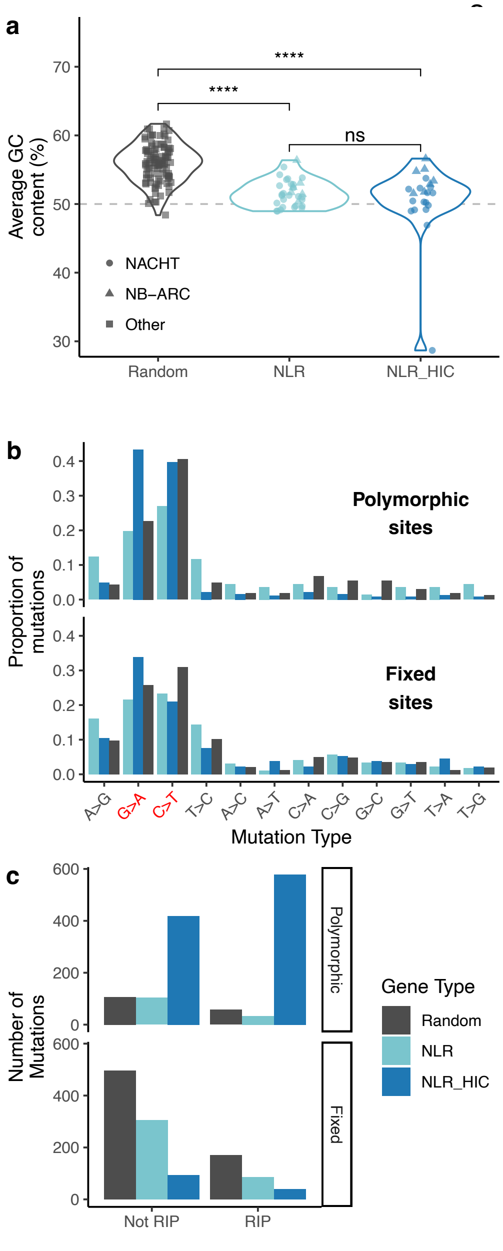


**Figure S1. Effect on repeat-induced point mutation (RIP) on *P. anserina* genes.** (**a**) Average GC content across gene classes. Significance determined with Kruskal-Wallis rank sum tests followed by a *post hoc* Dunn's test with Bonferroni correction (****: *p* <= 0.0001; ns: not significantly different). (**b**) Proportion of mutations, polymorphic or fixed within *P. anserina*, occurring in random genes, normal NLRs, or NLRs with HIC. The two transition types associated with RIP are highlighted in red. Mutation types are ordered such that transitions are first, followed by transversions. (**c**) Counts of mutations classified as either RIP-like (C>T mutation proceeded by an A or T, or G>A mutation preceeded by an A or T) or not RIP-like across gene types.

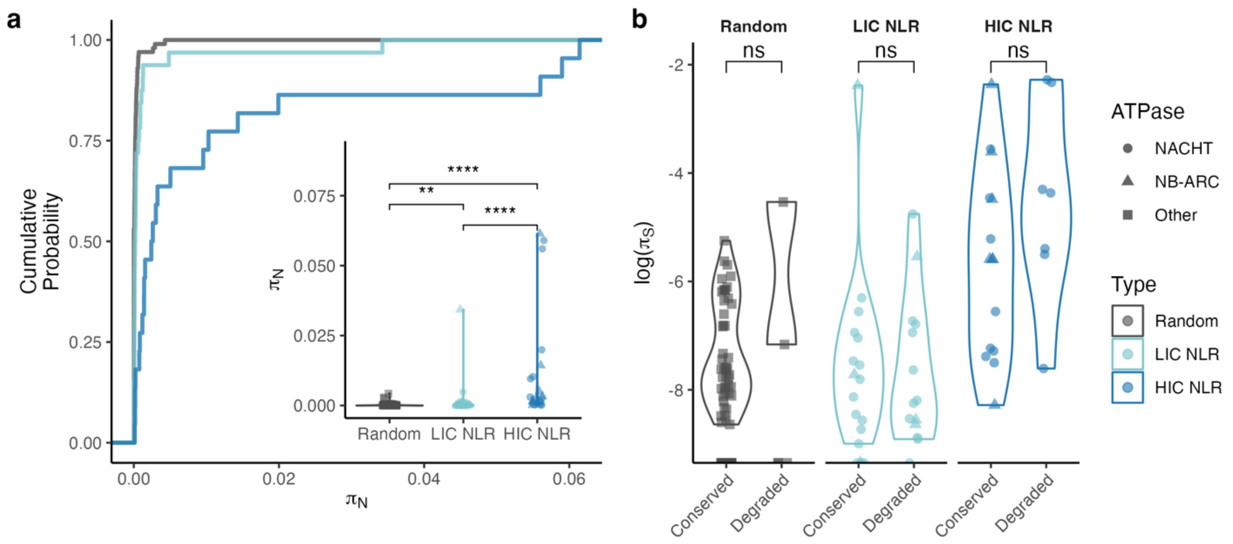


**Figure S2. Diversity metrics in *P. anserina* genes.** (**a**) Empirical cumulative distribution (ECDF) plot of pairwise-nucleotide non-synonymous diversity ($\pi_N$) for different gene classes, with distributions in a violin plot as inset. (**b**) Distribution of pairwise-nucleotide synonymous diversity ($\pi_S$) for different gene classes. Genes were further divided into "conserved" (putatively functional for all strains) and "degraded" (pseudogenized in all or some strains). Significance determined by Wilcoxon rank sum tests (ns: not significantly different).

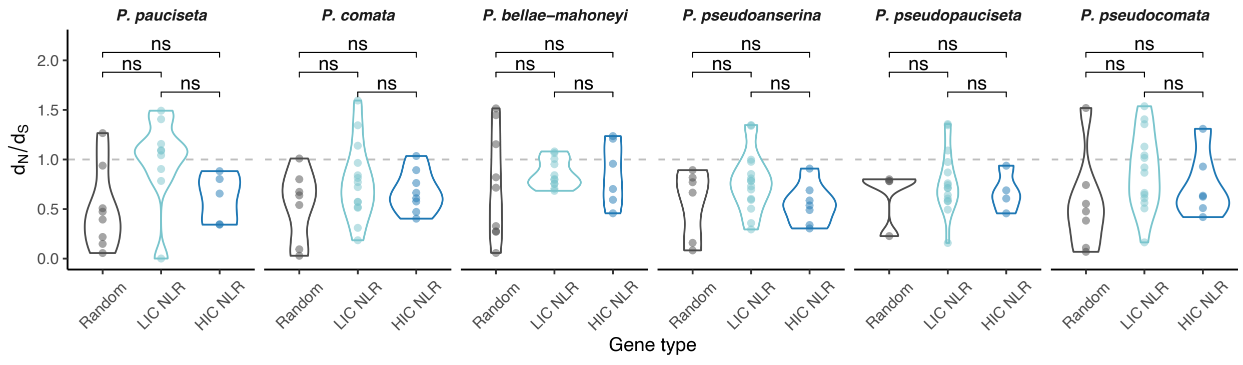


**Figure S3. Distribution of $d_N/d_S$ values between *P. anserina* (strain S+) and the ortholog of the other species (type strains) for degraded genes.** The dashed gray line marks $d_N = d_S$. Significance determined by Wilcoxon rank sum tests (ns: not significantly different).

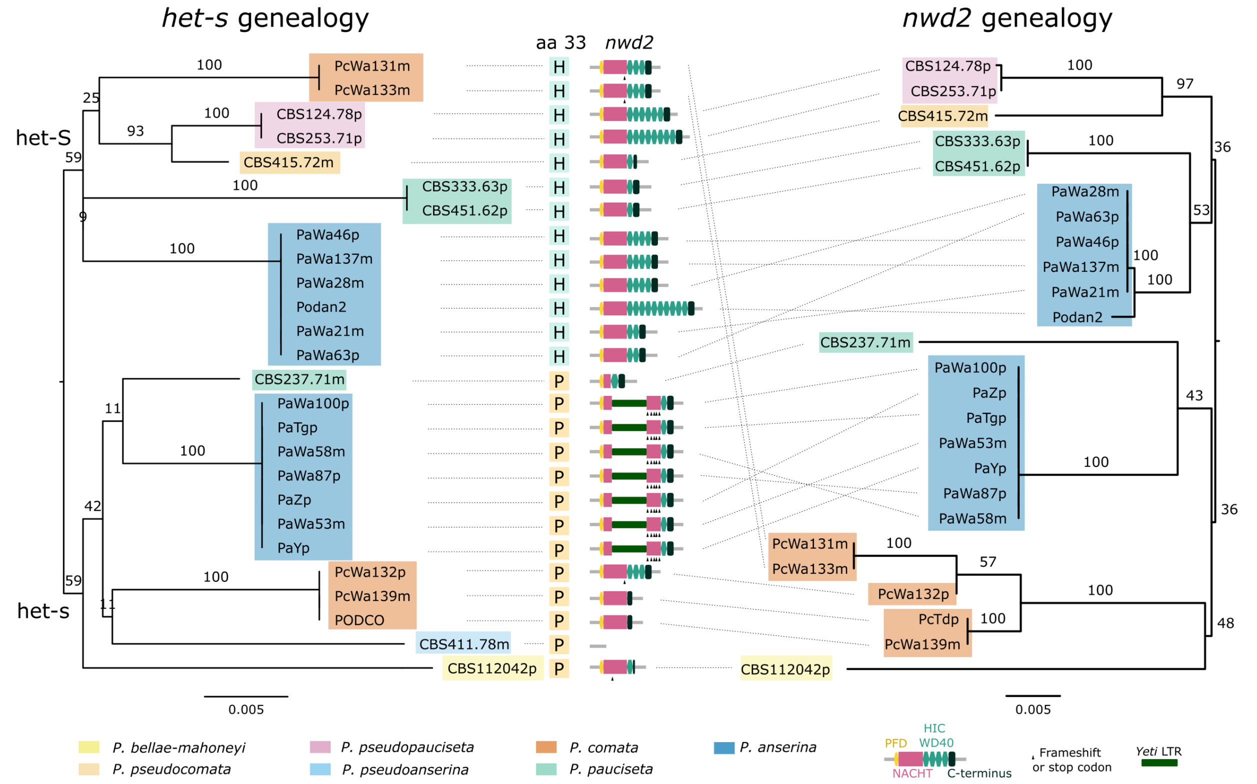


**Figure S4. Phylogenetic relationships of het-s and nwd2 in the *Podospora* species.** To the left, a maximum likelihood phylogeny of the *het-s* gene shows a split that matches the two alleles, which are defined by the presence of a proline (allele het-s) or a histidine (allele het-S) in the protein position 33. In the center of the figure, the corresponding position 33 of HET-s/S is illustrated along with a cartoon of the domain architecture of the linked *nwd2* allele. To the right, a maximum likelihood phylogeny of *nwd2* partially mirrors that of *het-s*. The exact number of WD40 repeats might be inaccurate for samples without long-read sequencing data. Branch support values correspond to standard non-parametric bootstrap (low within-species values are omitted for clarity). Branches are shown to scale as indicated by the scale bar (nucleotide substitutions per site). PFD: prion forming domain; NACHT: domain present in NAIO, CIITA, HET-E, TP-1; HIC WD40: repeats of the WD40 type with high internal conservation.

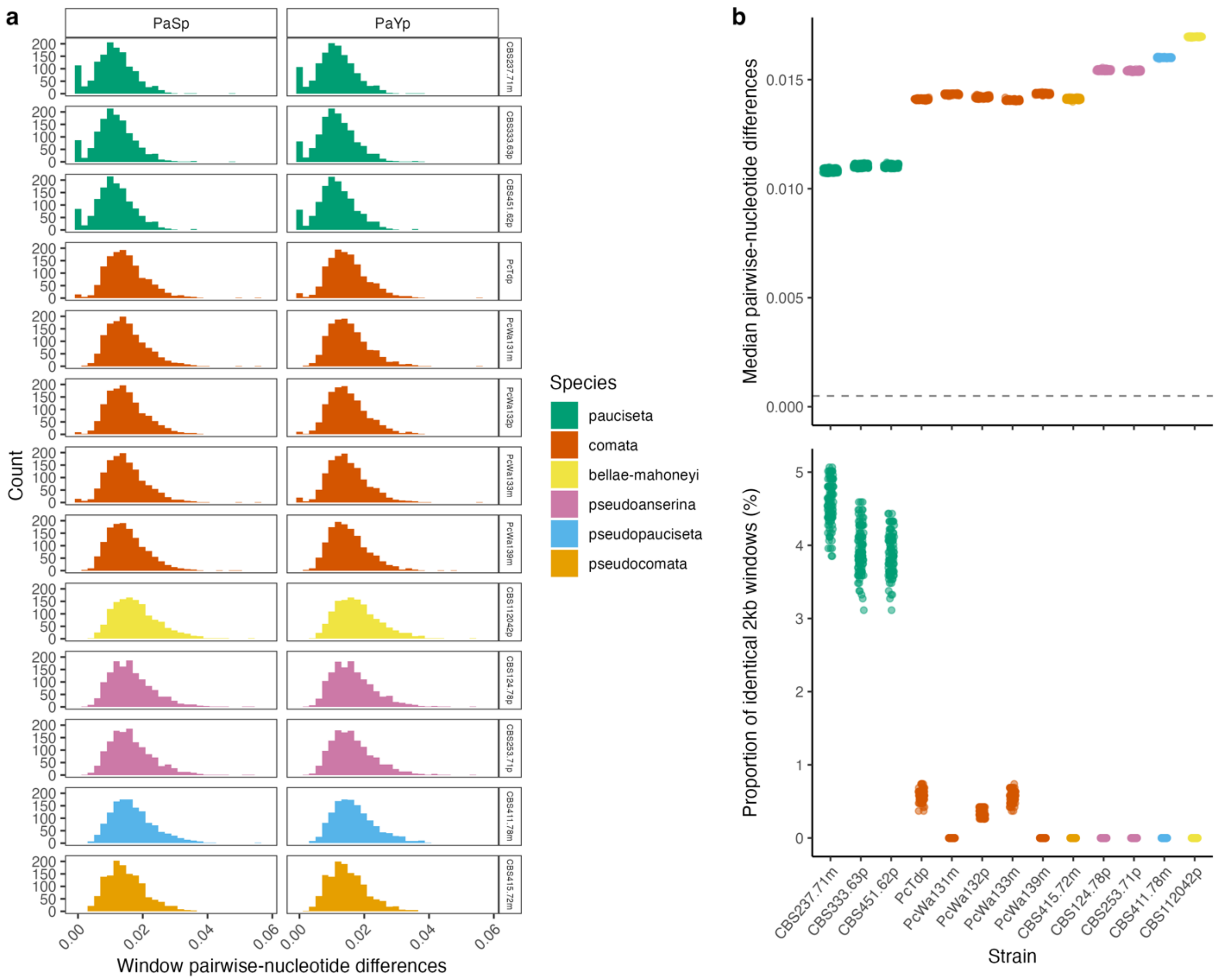


**Figure S5. Distribution of divergence between *Podospora* species in chromosome 4.** The pairwise-nucleotide differences (divergence) in 2 kb non-overlapping windows were calculated between two representative *P. anserina* strains (PaSp and PaYp) and each of the available strains of the other six species. In (**a**) the distribution of divergence values shows an excess of low divergence windows in comparisons with *P. pauciseta* and, to a lesser degree, some *P. comata* strains. The median window divergence values is very homogenous for all 106 *P. anserina* strains (**b**, top). However, the *Podospora* spp. strains have different proportions of nearly identical windows (divergence <= *P. anserina*'s π) relative to the 106 *P. anserina* strains, represented by individual data points (**b**, bottom). The dashed gray line marks the value of *P. anserina*'s π.

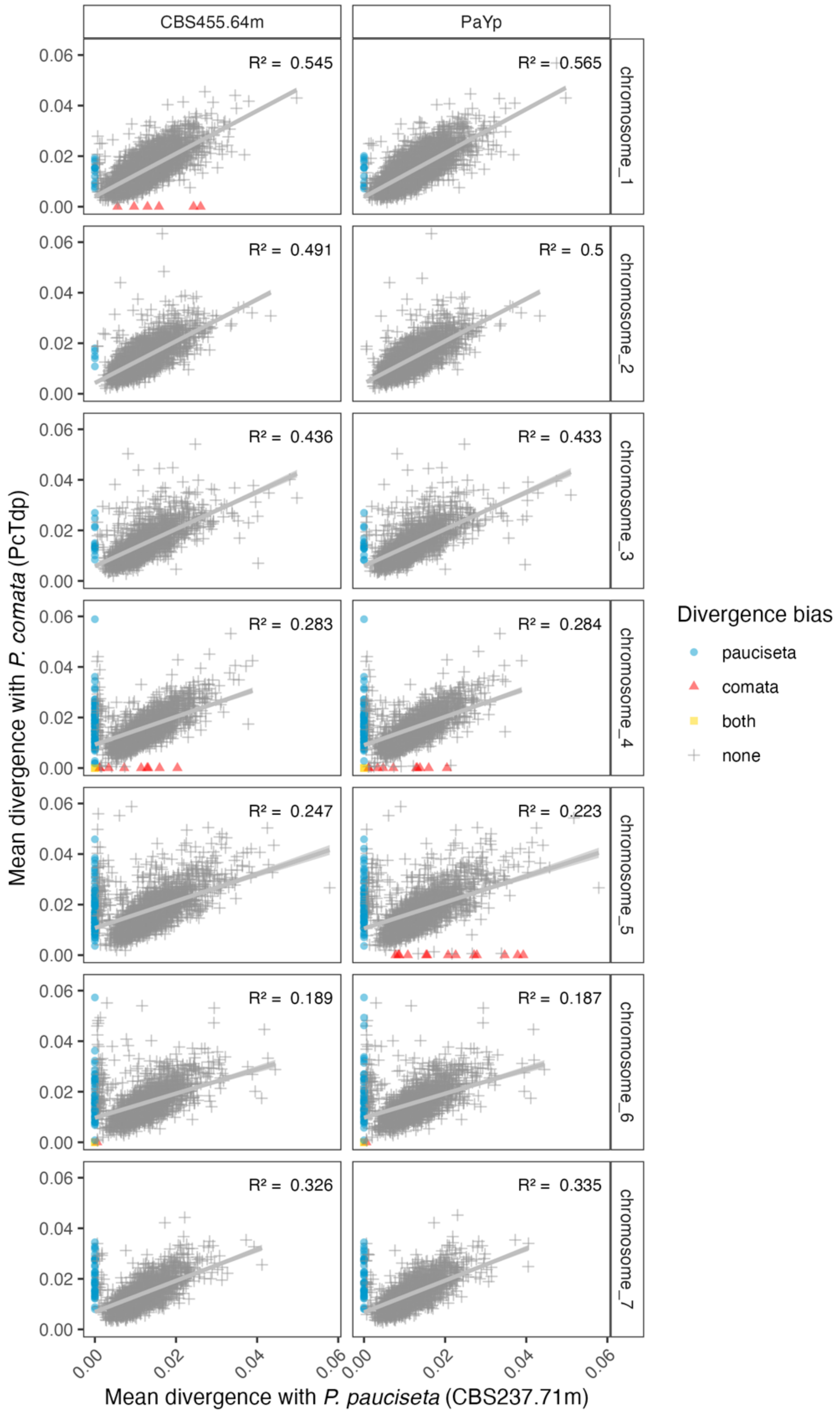


**Figure S6. The divergence between *P. anserina* and *P. pauciseta* is highly correlated with the divergence between *P. anserina* and *P. comata*.** Comparisons are shown for two example *P. anserina* strains (PaYp and CBS455.64m) against the type strain of either *P. pauciseta* (CBS237.71m) or *P.*

*comata* (PcTdp). Individual points correspond to nucleotide differences (divergence) in 2 kb non-overlapping windows (maximum 1 kb missing data), n = 14814 windows. The windows with divergence values equal or smaller than *P. anserina*'s intra-species diversity (π = 0.000497) are highlighted for either species or both. The $R^2$ values correspond to linear regressions.

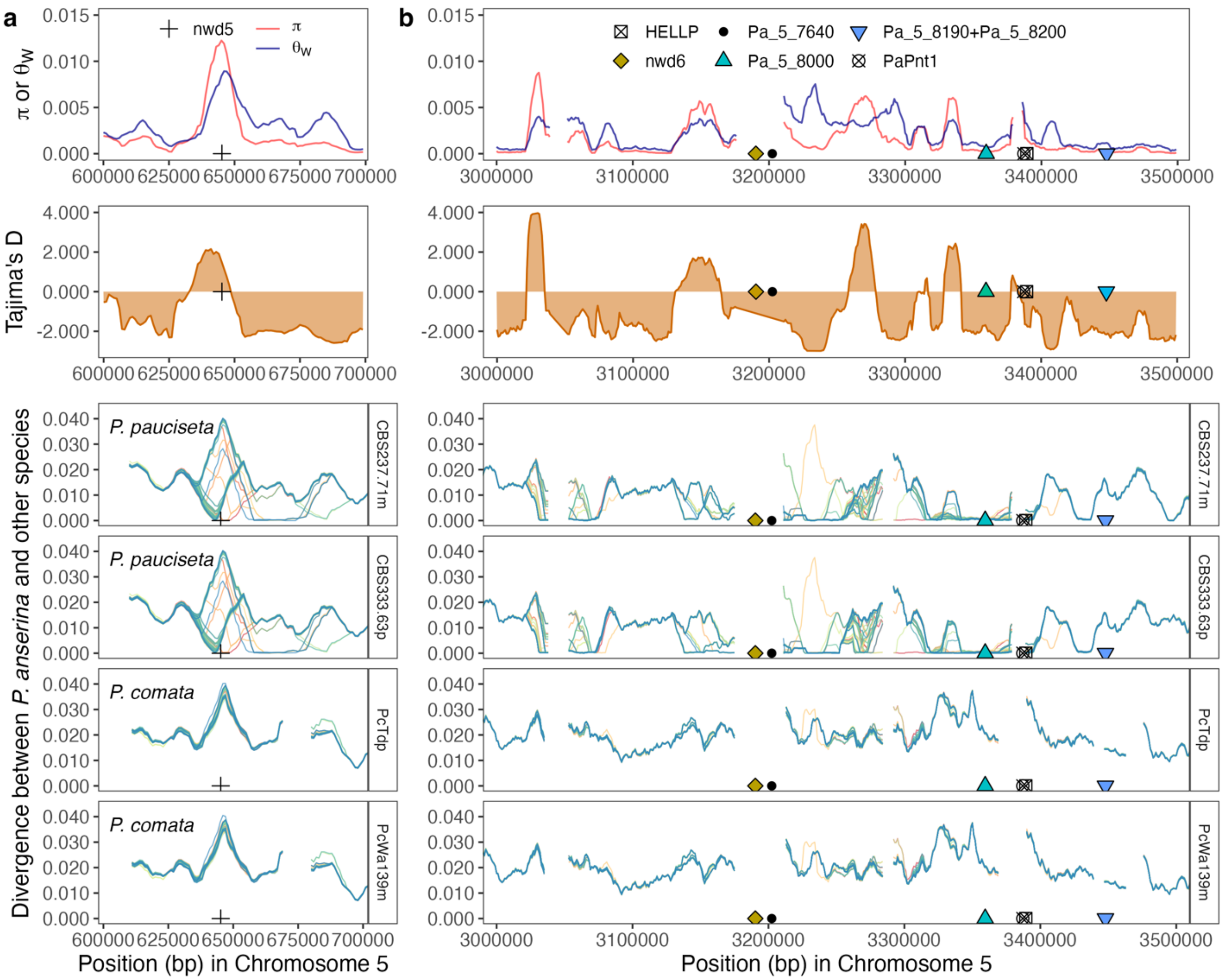


**Figure S7. Population diversity and sequence divergence of regions containing NLRs in chromosome 5.** The top panels correspond to the two metrics of genetic diversity π and Watterson's θ as well as Tajima's *D* in 10 kb windows (steps of 1 kb), as estimated in Ament-Velásquez et al. (2022) for the Wageningen population of *P. anserina*. The lower panels compare the pairwise-divergence between all 106 *P. anserina* strains vs. two different strains of either *P. pauciseta* and *P. comata*. Each *P. anserina* strain line was assigned a different color (gaps correspond to missing data). (a) Close-up to the *nwd5* region. (b) Larger complex region with several NLRs.

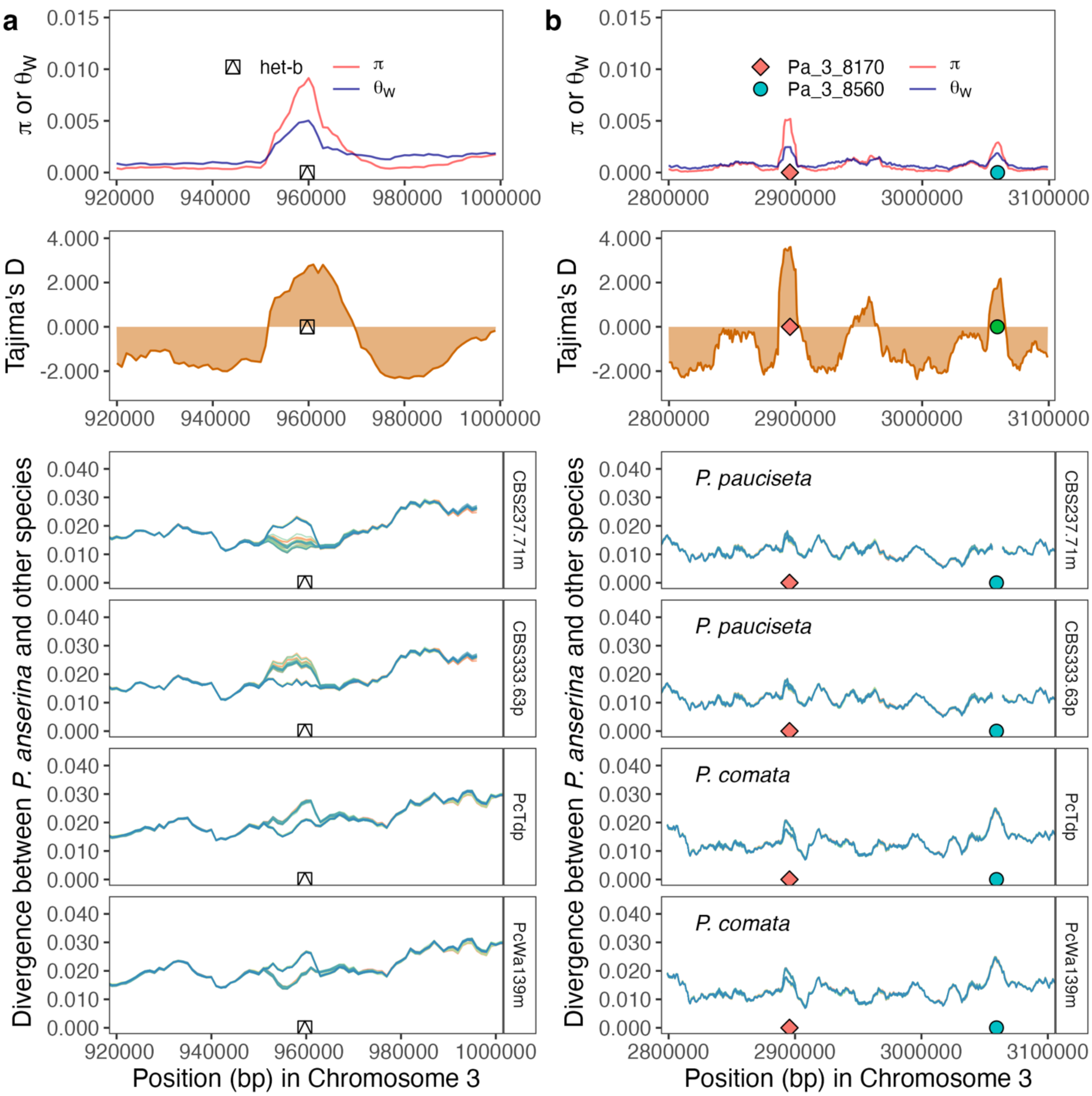


**Figure S8. Population diversity and sequence divergence around *het-b* and two NLRs potentially evolving under balancing selection.** The top panels correspond to the two metrics of genetic diversity π and Watterson's θ as well as Tajima's *D* in 10 kb windows (steps of 1 kb), as estimated in Ament-Velásquez et al. (2022) for the Wageningen population of *P. anserina*. The lower pannels compare the pairwise-divergence between all 106 *P. anserina* strains vs. two different strains of either *P. pauciseta* and *P. comata*. Each *P. anserina* strain line was assigned a different color. (a) Close-up to *het-b* region. (b) Close-up to *Pa_3_8170* and *Pa_3_8560* region. Notice that *Pa_3_8560* is absent in *P. pauciseta*;

conversely in *P. comata*, *Pa_3_8560* was in fact translocated to chromosome 5, but the reads mapped to its homolog in the *P. anserina* reference assembly (chromosome 3).

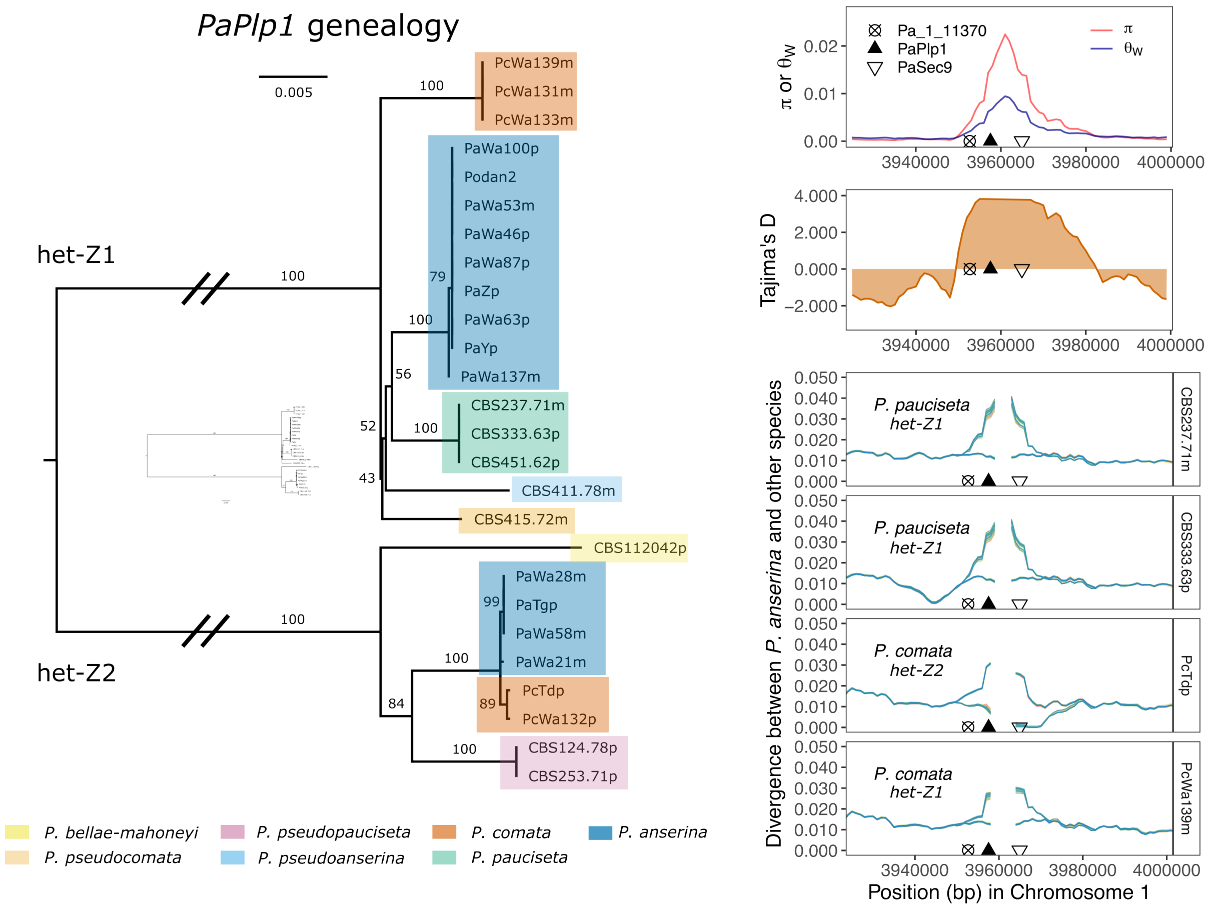


**Figure S9. Balancing selection and introgression patterns at *het-z*.** The *het-z* locus is conformed of the NLR *PaPlp1* and its cognate *PaSec9*. To the left, a maximum likelihood phylogeny of the *PaPlp1* gene shows a clear division of the two *het-z* alleles. Branches are shown to scale as indicated by the scale bar (nucleotide substitutions per site). Long internal branches have been reduced for clarity, with a miniature version of the full tree in the middle. To the right, the top panels correspond to the two metrics of genetic diversity π and Watterson's θ as well as Tajima's *D* in 10 kb windows (steps of 1 kb), as estimated in Ament-Velásquez et al. (2022) for the Wageningen population of *P. anserina*. The lower pannels compare the pairwise-divergence between all 106 *P. anserina* strains vs. two different strains of either *P. pauciseta* and *P. comata* (with their respective *het-z* alleles labeled). Each *P. anserina* strain line was assigned a different color (gaps correspond to missing data).

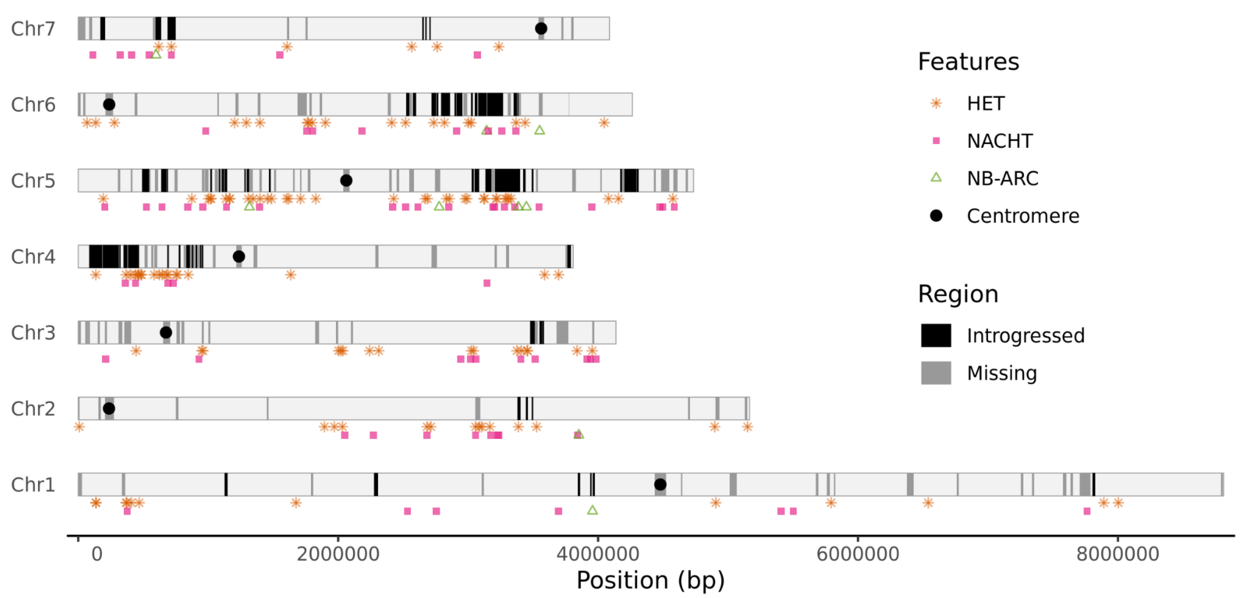


**Figure S10. Chromosomal distribution of some NLR-associated domains (as annotated by InterProScan) in relation to introgressed regions from *P. pauciseta* or *P. comata*.** The reference genome was divided in 10 Kpb-long overlapping windows (1 kb steps), which were classified as either introgressed (black) or regions with more than half missing data (gray).

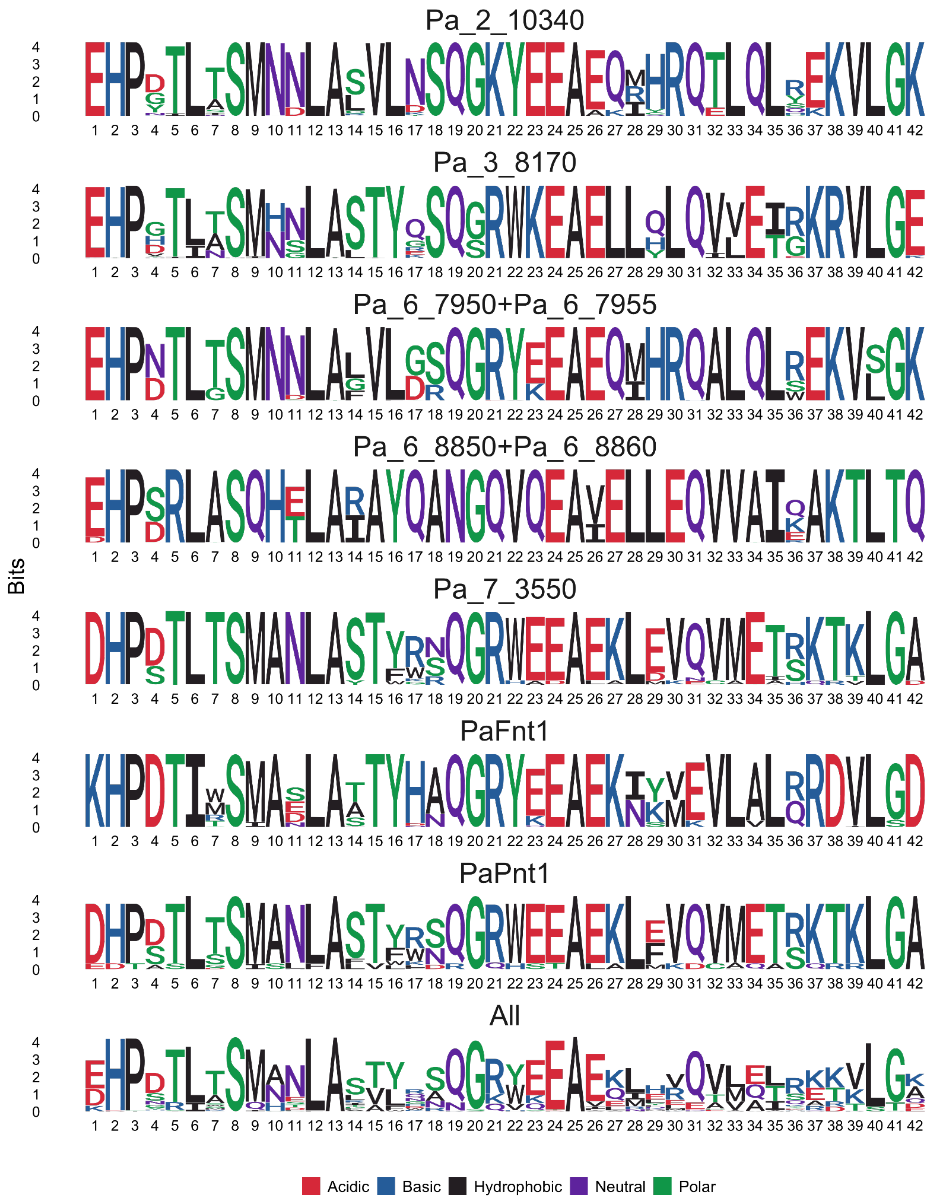


**Figure S11. TPR logo of the NB-ARC genes with HIC in *Podospora anserina*.**

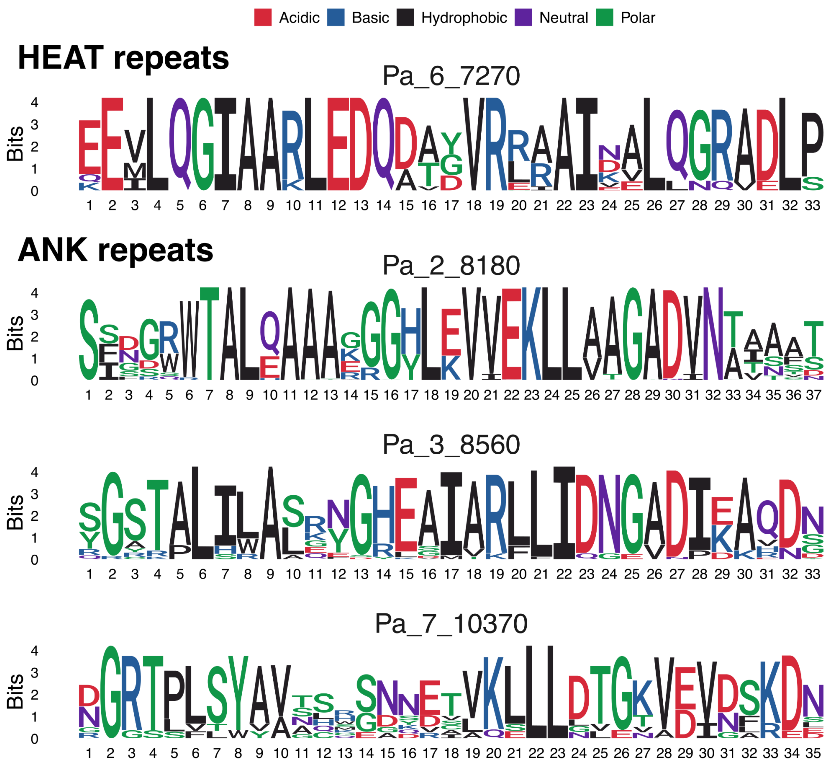


**Figure S12. Logos of HEAT and ANK NLR genes with HIC in *Podospora anserina*.**

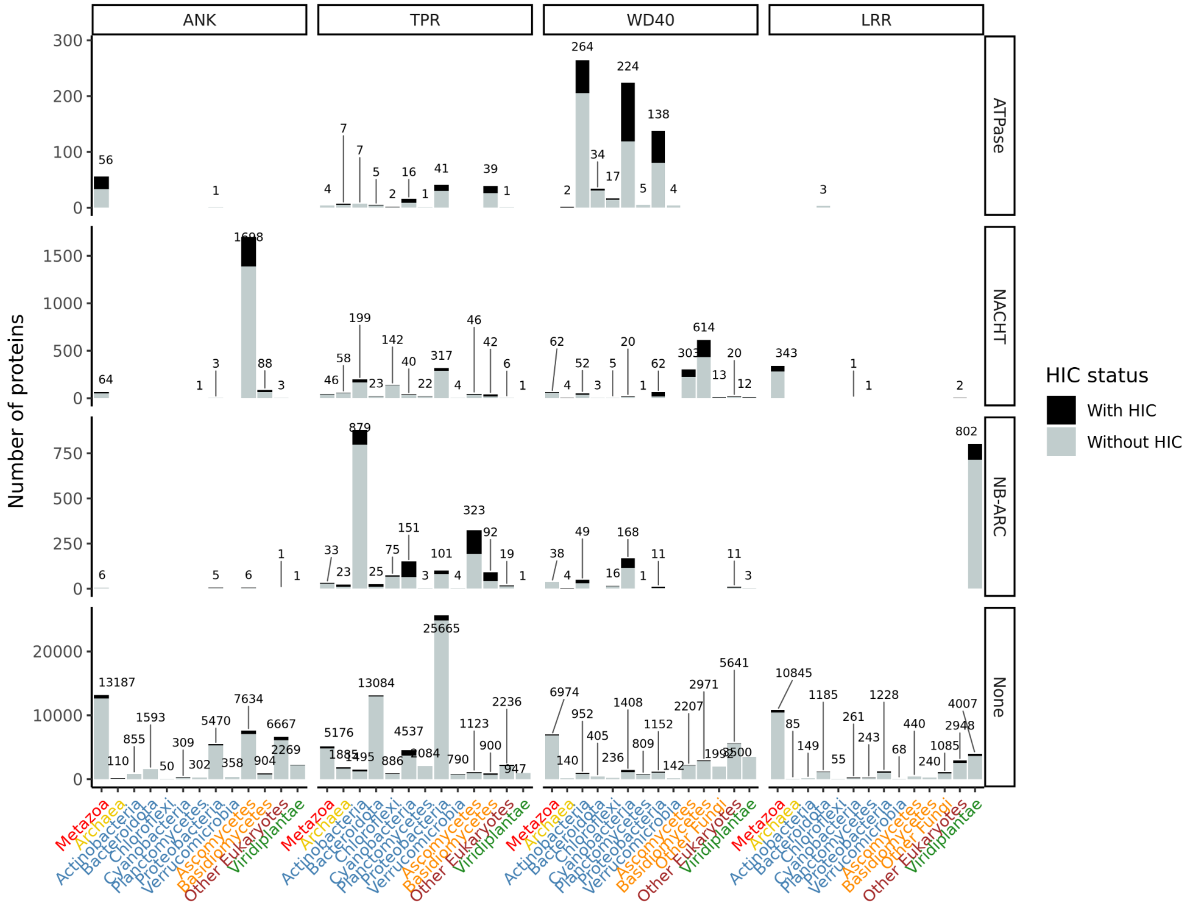


**Figure S13. Abundance of superstructure-forming repeat proteins from the SMART dataset per taxonomic group and repeat type after reducing redundancy.** The proportion of proteins with and without high internal conservation (HIC) is illustrated with gray and black colors. The “ATPase” category includes the nSTAND1 (PFAM code: PF20702), nSTAND2 (PF20703), and nSTAND3 (PF20720) domains. The NACHT category encompasses a number of related domains (PF13191, PF13238, PF13401, PF13479, PF13671, and PF24883). The NB-ARC category corresponds to the PFAM code PF00931. The “None” category are proteins that were not annotated with any of the above ATPase domains by InterProScan.

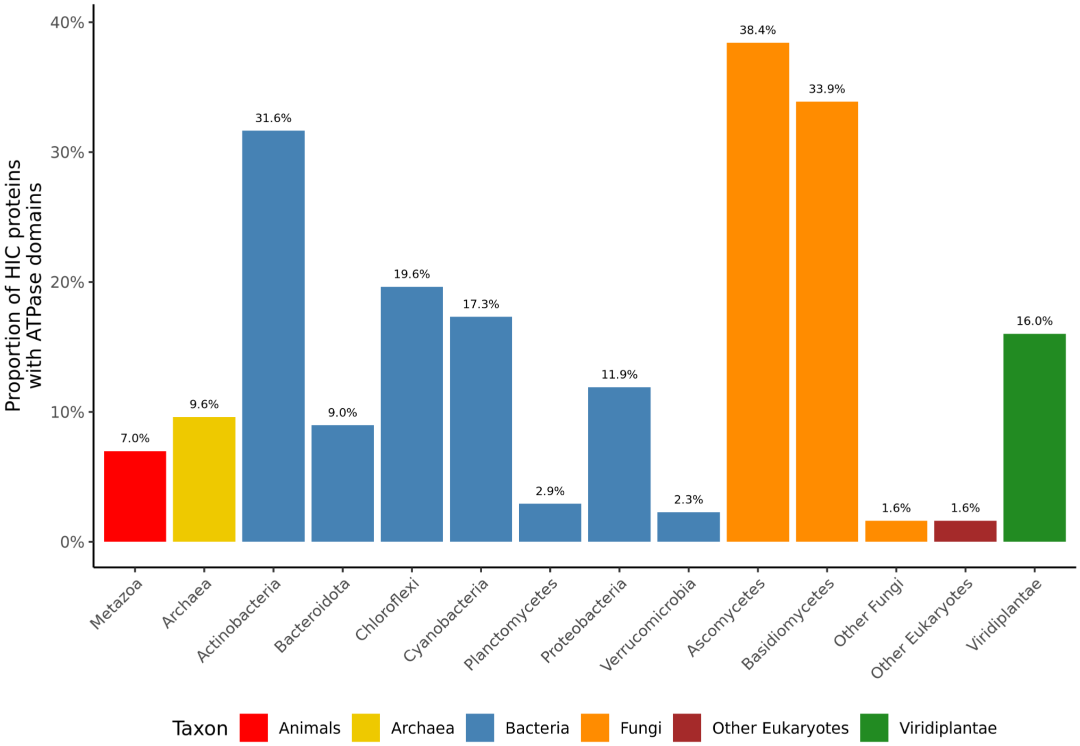


**Figure S14. Proportion of superstructure-forming repeat proteins with High Internal Conservation (HIC) and ATPase domains across various taxonomic groups.**
